# Modulation of habenula axon terminals supports action-outcome associations in larval zebrafish

**DOI:** 10.1101/2025.02.13.638047

**Authors:** Emanuele Paoli, Virginia Palieri, Amey Shenoy, Ruben Portugues

**Affiliations:** Institute of Neuroscience, Technical University of Munich, Germany; now at Champalimaud Center for the Unknown, Lisbon, Portugal; SyNergy Excellence Cluster, Munich, Germany; Bernstein Center for Computational Neuroscience, Munich, Germany; Max Planck Fellow Group - Mechanisms of Cognition, MPI Psychiatry, Munich, Germany; now at Department of Neurobiology and Behavior, Cornell University, Ithaca, NY, USA

## Abstract

Improving behavioral performance relies on the ability to associate decisions with their positive and negative outcomes. Although neurons that associate actions with their consequences have been identified across multiple brain regions, the circuit-level mechanisms underlying this integration remain poorly understood.

Here, using an operant thermoregulatory assay, we show that larval zebrafish maintain a short-term memory of action-outcome associations and that the dorsal habenula-interpeduncular nucleus (dHb–IPN) pathway is necessary for this process. Consistently, a population of intermediate IPN neurons encodes actions only when they lead to a thermal reward, suggesting a major role in establishing short-term associations and influencing subsequent decisions. We then combine circuit mapping and axon imaging to show that actions and reward signals are conveyed by GABAergic prepontine and glutamatergic dHb neurons, respectively. Finally, the integration between motor and sensory streams relies on presynaptic GABA_B_ receptor–mediated modulation of dHb axon terminals by prepontine neurons.

These results link a crucial computation for adaptive behavior to a specific circuit mechanism.

## Introduction

The ability to associate actions with their outcomes forms the basis for effective interactions with our environment, enabling animals to adapt their behavior based on past experience^1^. At its fundamental level, this process first requires the brain to detect which action led to a positive or negative event. For example, neurons can encode specific actions only when they are followed by a reward^2^. In this way, downstream circuits are informed about an action’s effectiveness in meeting the animal’s needs and can internally maintain this information for future use. Although neurons implementing such action-outcome code have been found in diverse brain regions^2–5^, a general circuit principle explaining how this code emerges and its relevance for behavior remains elusive.

Here, we studied how actions and reward signals are integrated in the highly structured and evolutionary conserved dorsal habenula (dHb)-interpeduncular nucleus (IPN) pathway in larval zebrafish^6,7^.

Using a thermoregulatory operant assay, we found that zebrafish maintain short-term associations between their actions and thermal reward, to adjust their reorientation behavior and minimize exposure to cold temperatures^8–14^. Delaying the reward from the motor event impairs performance, indicating that associations rely on the precise timing between a successful choice and its sensory outcome.

We next found that the dHb–IPN pathway is necessary for establishing these associations. A population of neurons in the intermediate IPN (iIPN) encodes actions only when they are followed by a thermal reward within a short time window, effectively multiplying motor and sensory streams^15^. These neurons are organized in two lateralized clusters that send their dendrites into well defined neuropil regions called glomeruli. Neurons in the right iIPN glomerulus selectively multiply counter-clockwise turnings and reward while the left glomerulus has the opposite action tuning. This integration is temporally precise, with a time window similar to the one found in the operant assay.

Using a combination of electron microscopy, photoactivations and functional imaging we reconstructed the entire circuit upstream of iIPN neurons, alongside their neurotransmitter identity and functional profile. We found that the glutamatergic right subnucleus of the dHb broadcasts thermal reward information to both the left and right iIPN glomerulus. However, this input is locally modulated at the dHb axon terminals depending on the preceding behavioral choice. Therefore, here we propose that the integration of actions and outcomes arises at the presynaptic terminals of reward neurons through local modulation rather than postsynaptically. Finally, dHb axon modulation relies on a non-canonical GABA_B_ receptor–mediated potentiation of dHb glutamate release and is driven by a lateralized GABAergic population of prepontine neurons conveying choice signals.

## Results

### Larval zebrafish maintain short-term action-outcome associations to drive adaptive thermoregulatory behavior

Zebrafish larvae seek to maintain a stable body temperature of *∼* 25.3 °C^16^. In natural settings, this equilibrium is achieved by monitoring temperature changes detected during swims and by reacting to deviations from this set-point.

Here, we decided to study how associations between actions and outcomes are formed in the framework of temperature regulation. More specifically, whether fish is able to associate and remember specific swim types with the outcome of warming its body in a cold environment (thermal reward). To this end, we developed an assay in which rapid body temperature changes can be experimentally controlled and delivered on a swim-by-swim basis.

Fish could freely swim in a chamber with water at 20 °C. Their position was continuously monitored with a high-speed camera, and a set of galvanometric mirrors was used to direct an infrared laser onto the fish’s body (Figure S1a, see Methods)^17^. The laser was calibrated to increase body temperature by *∼* 1 °C (Figure S1b-d, thermal reward). Given zebrafish setpoint, this condition is expected to have a positive valence for the larvae and to be preferred over the opposite condition (Figure 1a).

**Figure 1.**
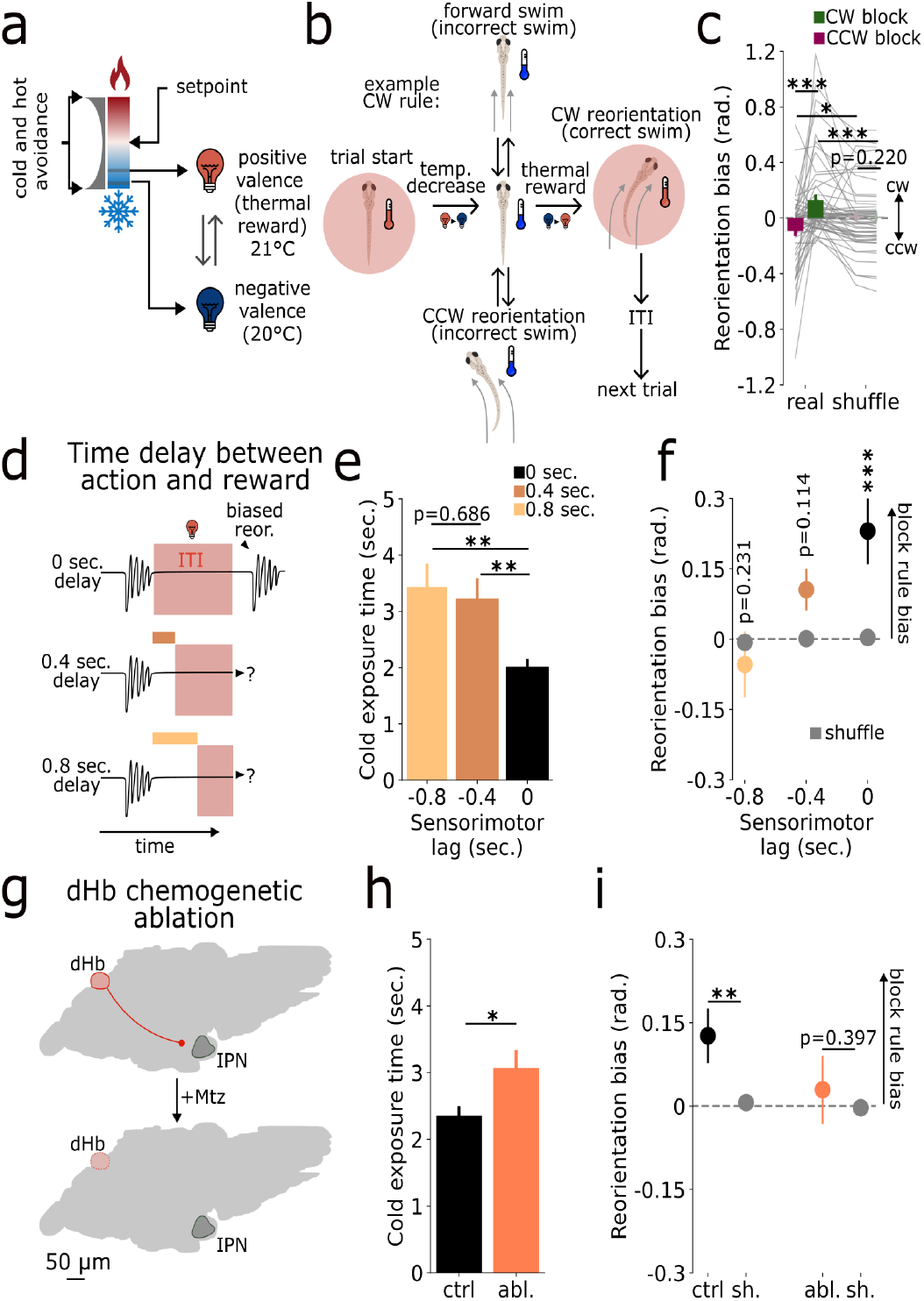
The dHb support the maintenance of short-term action-outcome associations. **(a)** Experimental concept. Thermoregulation is a symmetric process centered around the animal’s setpoint. Deviations from this value trigger cold or hot avoidance behaviors. In a cold environment, the IR laser heating will have a positive valence for the animal. **(b)** Trial structure of the operant assay. Fish can increase body temperature by turning in the rule direction (either CCW or CW). **(c)** First swim average reorientation (mean ± s.e.m.) after temperature decrease for CCW (magenta) and CW (green) blocks (first two bars) and their shuffles (third and fourth bars). Each line represents a single fish. **(d)** Concept of the action-outcome delay experiment. **(e)** Cold exposure time corrected for delay condition (mean ± s.e.m.). Each bar represents the time between temperature decrease and the correct swim (black: 0 sec. delay, dark brown: 0.4 sec. delay, light brown: 0.8 sec. delay). **(f)** First swim average reorientation (mean ± s.e.m.) at temperature decrease, with CCW and CW blocks pooled together (black: 0 sec. delay, dark brown: 0.4 sec. delay, light brown: 0.8 sec. delay, gray: shuffles). **(g)** Schematic of dHb chemogenetic ablation experiment. **(h)** Cold exposure time (mean ± s.e.m.). Each bar represents the time between temperature decrease and the correct swim (black: treatment control group, pink: ablation group, gray: shuffles). **(i)** First swim average reorientation (mean ± s.e.m.) at temperature decrease, with CCW and CW blocks pooled together (black: treatment control group, pink: ablation group, gray: shuffles).

We first tested this preference using an assay where laser stimulation was delivered contingent on the fish entering a rectangular region of interest (ROI) within the arena (Figure S1e). Importantly, no positional cues indicated to the fish where this ROI was.

As expected, larvae (n = 21 animals) consistently preferred the warmer region compared to a control group (n = 5 animals) without IR heating (Figure S1f-h).

We next designed an operant thermoregulatory assay in which thermal reward is paired with either clockwise (CW) or counterclockwise (CCW) turns^8–14^.

In this paradigm fish could increase their body temperature by turning in a specific direction (rule direction) and consequently minimize the time spent at the colder temperature. The experiment (n = 54 animals) consisted of 60 trials: 30 conditioned in one swim turn direction and 30 in the opposite. Each trial (Figure 1b) began with a period (inter-trial interval or ITI) of at least 5 seconds (see later) during which an IR laser followed the fish. After this the laser was turned off, initiating a cold period of up to 10 seconds. During this cold period, turning in the rule direction allowed the fish to rapidly increase their body temperature again by receiving the thermal reward. This would lead to the next ITI before starting the next trial. If the fish did not turn in the rule direction within 10 seconds, the trial was aborted. To ensure that the fish had space to turn freely in both directions, trials were initiated only when the animal was distant from the arena borders (see Methods), resulting in a distribution of ITIs (Figure S2a).

We reasoned that if zebrafish associate actions and thermal reward, their first swim during the cold period in each block should be biased toward rule direction. Indeed, the average angular velocity and cumulative reorientation of the first swim confirmed this hypothesis (Figure 1c and Figure S2b–c). Directional bias was not observed during the ITIs (Figure S2d–e), indicating that the conditioned response is driven by the aversive temperature drop. Moreover, it was only detectable following short ITIs (< 7.5 seconds; Figure S2f), suggesting that action-outcome associations in this paradigm rely on a short-term memory mechanism (Figure S2g).

We next assessed whether the precise timing between the correct swim and thermal reward is critical for short-term associations.

To test this, we introduced a delay of 0, 0.4, or 0.8 seconds between the two events, and repeated the assay (Figure 1d; n = 20 animals per condition). Indeed, we observed a significant impairment in task performance, as reflected by the average cold exposure time (corrected for delay; Figure 1e) and reorientation bias (Figure 1f).

### Short-term action–outcome associations require dHb function

Previous studies have shown that the zebrafish dorsal habenula (dHb) is crucial for experience-dependent behaviors^18–21^ and thermal gradient navigation^16^. To test whether this structure is required in our assay, we performed chemogenetic ablation of dHb neurons (Figure 1g, Figure S3a; see Methods) and repeated the behavioral experiment (ablation group: n = 71, treatment control group: n = 72). In accordance with previous findings, we found that ablation significantly impaired cold exposure time (Figure 1h) and reduced reorientation bias (Figure 1i), suggesting a deficit in establishing action-outcome associations.

Previous research has shown that, when exposed to homeostatic threats, zebrafish generate large-amplitude turns as a defensive behavioral strategy^16,22–24^. Therefore, we reasoned that if dHb ablation led to a general deficit in sensory processing, then this innate behavior should be affected as well. However, the reorientation magnitude (regardless of CCW or CW direction) of the same swims quantified earlier (first swim after temperature decrease) was similar in the two experimental groups and higher than shuffled controls (Figure S3b). We also found no evidence of motor deficits, based on the analysis of multiple swimming kinematic parameters (Figure S3c-f).

Together, these results suggest that the dHb is not directly involved in motor control or innate aversive sensorimotor responses. Instead, its role appears to be specific to guide behavior based on past sensorimotor experience^16^.

### Thermal reward responses in r-dHb neurons

Once established a crucial role for the dHb, we aimed at studying how movements and reward signals are encoded and possibly integrated in this region and its unique downstream target, the interpeduncular nucleus (IPN, Figure 2aI, Figure S4a-d). To this end, we developed a head-fixed protocol where temperature changes were temporally locked to the end of the animal’s movements, which were detected online throughout the experiment (Figure 2aII, see Methods). To facilitate statistical comparisons across sensorimotor conditions, each swim (regardless of amplitude or direction) had a probability of switching the current laser state from on to off or vice versa. As a result, swim decisions can be categorized by combinations of swim type (CCW, CW or forward) and outcomes (thermal reward, temperature decrease and no temperature change) (Figure 2aIII, (Figure S4e).

**Figure 2.**
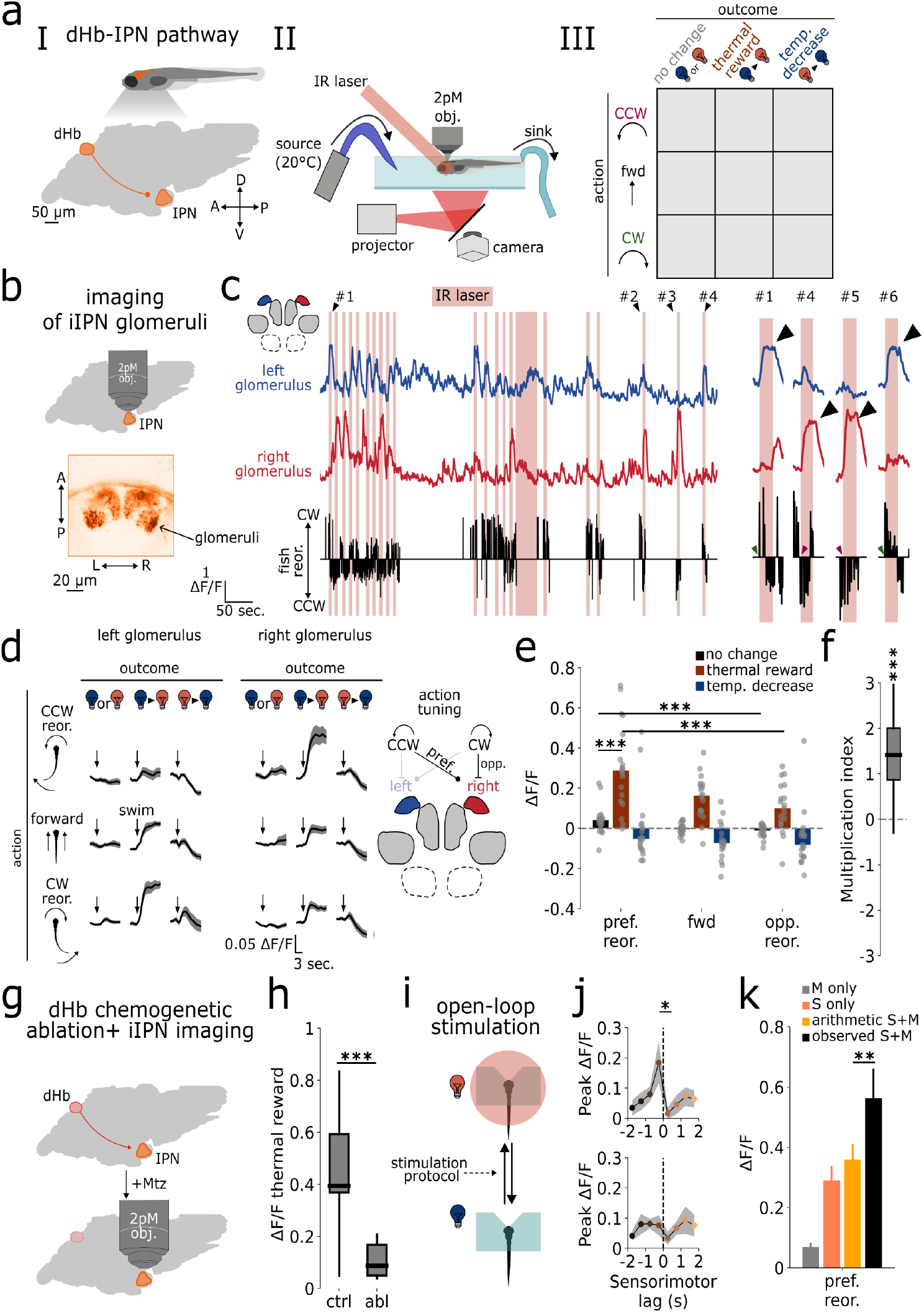
Emergence of action-outcome coding in iIPN neurons. **(a)** I: Schematic of the dHb-IPN pathway. II: Head-fixed preparation for imaging experiments. III: Protocol structure: Temperature changes are stochastically triggered at swim termination, allowing the evaluation of multiple action-outcome combinations. **(b)** Top: schematic of 2-photon calcium imaging experiments in the dendrites of iIPN neurons (glomeruli). Bottom: Example horizontal section of the iIPN glomeruli. **c)** Left: example traces from the left (blue) and right (red) glomerulus. The black trace below represents the animal’s behavior, with upward and downward spikes indicating CW and CCW turns, respectively. Green and magenta arrowheads point to CW+thermal reward and CCW+thermal reward events, respectively. Right: magnified action-outcome events along with glomerular activity. **(d)** Right- and left-glomerulus triggered average (mean ± s.e.m.) for different action-outcome combinations. **(e)** Left: anatomical-based quantification. Blue: temperature decrease. Brown: thermal reward. Black: no temperature change. pref. reor.: contralateral reorientation relative to the glomerulus’ anatomical position. fwd: forward swim. opp. reor.: ipsilateral reorientation relative to the glomerulus’ anatomical position. Each dot represents a single fish, with bars indicating the median across fish. Right: Schematic of glomerulus action-tuning. **(f)** Multiplication index. **(g)** Schematic of dHb ablation combined with iIPN 2-photon calcium imaging. **(h)** Average ΔF/F increase at thermal reward delivery for controls and ablation group. **(i)** Schematic of open-loop experiment. **(j)** Peak ΔF/F (mean ± s.e.m.) for different sensorimotor lags. Colors depict different sensorimotor lag bins. Lags are color-coded from black (negative lag, choice precedes outcome) to yellow (positive lag, opposite scenario). Left: Triggered average for different bins across fish. **(k)** ΔF/F (mean ± s.e.m.) responses for the preferred turn direction that was not followed by thermal reward (motor only or M only, gray bar), thermal reward that was not preceded by any swim (sensory only or S only, pink bar), arithmetic addition of the two (sensory + motor or arithmetic S + M, yellow bar) and the observed responses when preferred turns are followed within 1 second by the thermal reward (observed sensory + motor or observed S + M, black bar).

We started by imaging neurons in the right subnucleus of the dHb (r-dHb, n = 14 animals), known to contain temperature-sensitive neurons^16,22^(Figure S5a). Indeed, we found neurons tuned to both thermal reward and the opposite temperature change (34.5%, 16.9% of the total recorded neurons, Figure S5b-d, see Methods). Thermal reward neurons had heterogeneous response profiles, which we categorized as fast-adapting (transient increase with rapid adaptation), fast non-adapting (sustained increase), and slow (Figure S5e-g).

### iIPN neurons encode action identity contingent on thermal reward

We then imaged the intermediate portion of the IPN (iIPN), a target subnucleus of the r-dHb. Here, local neurons cluster their dendrites into well-defined regions called glomeruli^25^. To study their activity, we used a transgenic line that labels a population of excitatory (vglut2a+) neurons and recorded from three bilateral glomeruli (Figure 2b, Figure S6a–b; n = 19 animals).

To our surprise, only one glomerulus per side showed elevated activity during our protocol (from now on referred to simply as the glomerulus). In stark contrast to what we observed in the r-dHb, glomerular activity was variable, with clear transients during thermal reward occurring only a fraction of the time. Closer inspection revealed a strong relationship between this variability and the animal’s behavior. Figure 2c provides an example, showing the activity of the left (blue trace) and right glomerulus (red trace). When the animal turns CCW or CW but this swim does not lead to a temperature change (no outcome), the glomeruli are almost silent. However, when the same type of movement leads to the thermal reward, the glomerulus located in the hemisphere contralateral to the turned direction strongly increases its fluorescence (Figure 2d, Figure S6c). Note that the change in temperature is delivered at movement termination and, considering the stochastic nature of our stimulation protocol, the animal cannot predict the outcome of each swim. Interestingly, once active, glomerular activity does not return to baseline like motor-modulated cells, but instead remains active throughout the stimulation period (Figure 2c right, black arrows).

We then quantified the interaction between different action-outcome events on the basis of their hemispheric position. First, we computed the evoked ΔF/F based on all combinations of actions and outcomes. Then, we expressed each glomerulus’ tuning relative to its position and swim direction. In this analysis, the “preferred reorientation” (pref. reor.) condition includes responses from the left glomerulus during CW swims and the right glomerulus during CCW swims, categorized by different outcomes. Conversely, the “opposite reorientation” (opp. reor.) condition groups responses from left glomerulus during CCW swims and right glomerulus during CW swims. This analysis revealed clear anatomical structuring (Figure 2e) suggesting an underlying lateralized circuit architecture.

The difference in evoked activity between opposite reorientations (CCW and CW) with and without thermal reward, suggests that these neurons multiply specific motor actions and thermal feedback. To test whether this observation is a general feature in our dataset, we designed an index (multiplication index or MI, see Methods) specifically tailored to test this form of multiplicative responses. An MI close to zero indicates a canonical sensory-like neuron (responses to temperature increase without action tuning) or an additive interaction between the sensory and motor variables. A positive MI indicates multiplicative interaction, whereas a negative MI suggests directional swim modulation only in absence of thermal feedback. This analysis revealed an MI of 1.413, corresponding to a *∼* 4-fold increase in activity when the preferred action was followed by reward (Figure 2f).

We next imaged population activity in the somata of iIPN neurons (Figure S7a, n = 17 animals) and compared it to the activity measured in the upstream r-dHb. The MI distribution of the latter was centered at zero, suggesting, in line with previous results, sensory-like responses. In contrast, iIPN neurons were shifted toward positive values (Figure S6b-c). Finally, we tested whether iIPN thermal reward modulation depends on an intact dHb. We repeated the dHb chemogenetic ablation and after 1 day of recovery, we imaged iIPN neurons (Figure 2g). Compared to controls, the ablation group significantly reduced the magnitude of evoked ΔF/F at reward delivery (Figure 2h), suggesting that the r-dHb is necessary for the sensory outcome component of iIPN responses.

### Integration of actions and thermal reward in iIPN neurons is temporally restricted and asymmetric

To determine how sensory and motor signals are integrated in these neurons, we examined neuronal activity during an open-loop protocol, in which sensation and action are decoupled (Figure 2i, n = 21 animals, see Methods).

Even under these conditions, some stimulation trials evoked clear calcium transients, especially when the fish swam close to thermal reward (Figure S7d). We next calculated the evoked ΔF/F as a function of the lag between the two event types (sensorimotor lag). Negative lag values indicated that reward delivery followed the swim, while positive values indicated the opposite sequence. The absolute value of the lag corresponded to the interval between the two events (see Methods, Figure S7e, left). A neuron acting as a coincidence detector would show a symmetric response profile around zero lag, whereas a sequence detector would display an asymmetric response (Figure S7e right). Strikingly, the two glomeruli strongly responded when the preferred motor event preceded the sensory event by less than one second (negative lags, Figure 2j, top: preferred reorientation, bottom: non-preferred reorientation) but not for the opposite sequence. Finally, we evaluated glomerular responses during the preferred swim direction closely followed by thermal reward (< 1 second) and compared it with the arithmetic sum of the two events in isolation (see Methods). We found that addition alone cannot account for glomerular responses (Figure 2k), further supporting the multiplication hypothesis.

Overall, our imaging experiments revealed a significant transformation between r-dHb and iIPN neurons. While r-dHb activity signals thermal reward, the iIPN encodes which action successfully led to that event by multiplying motor and sensory variables. Furthermore, this action–outcome coding is localized to a specific iIPN glomerulus. This, together with the lateralized action tuning, strongly suggests a well-defined circuit underlying this computation.

### iIPN neurons receive habenular and prepontine inputs

To elucidate the circuit organization underlying iIPN activity, we aimed at reconstructing these neurons along with neurons targeting the glomerulus using a SBEM dataset^26^. We reconstructed 41 right-iIPN neurons whose dendrite was located within the IPN neuropil (Figure 3a). They had a bipolar morphology, a single dendritic tree and their axons reached a neuropil region near the dorsal raphe nucleus (DRN) (Figure 3b first column, Figure S8a). Although these neurons projected outside the IPN, they remained close, still within rhombomere 1/2.

**Figure 3.**
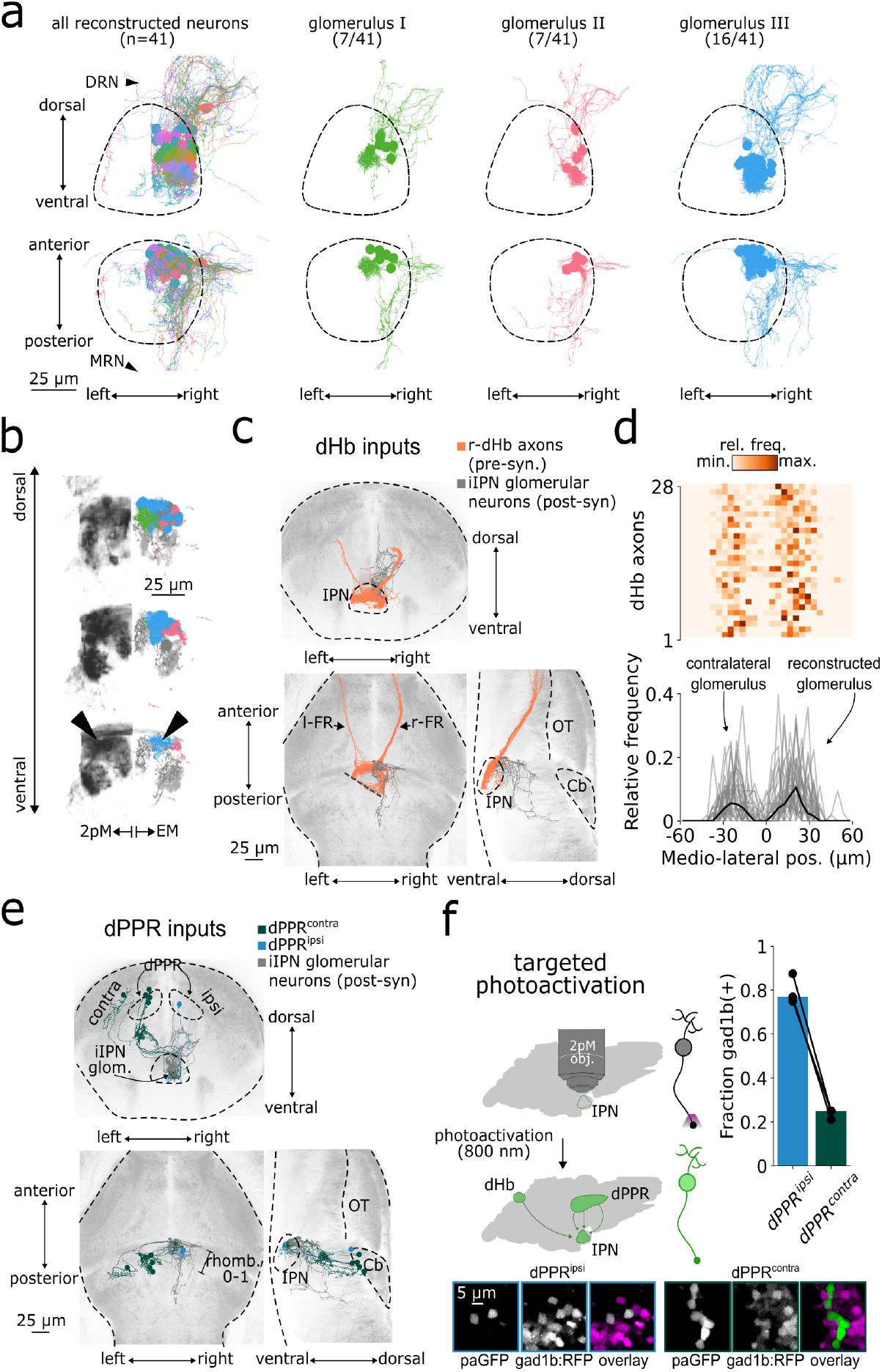
iIPN circuit organization. **(a)** iIPN reconstructions. Columns represent all the reconstructed neurons (first column, color randomly assigned, DRN: dorsal raphe nucleus, MRN: median raphe nucleus) and the three identified glomeruli (green: glomerulus I, pink: glomerulus II, blue: glomerulus III). Frontal projection on the top row and horizontal projection on the bottom row. Dashed regions enclose the IPN neuropil. Reference space from mapZebrain atlas. **(b)** Co-localization of EM and functional imaging dataset. Left: transgenic line labeling iIPN neurons. Right: EM reconstructions with the same color code as in (a). Arrowhead point to the same positions in the two modalities. Reference space from mapZebrain atlas. **(c)** dHb axons (pink) reconstructed as presynaptic partners of seed iIPN neurons (gray). Top: frontal projection. Left bottom: horizontal projection. Right bottom: sagittal projection. The dashed line in the horizontal projection indicatesa dataset discontinuity where most of the dHb axons were missed (OT: optic tectum, Cb: cerebellum, l-FR: left fasciculus retroflexus, r-FR: right fasciculus retroflexus). Reference space and anatomy from mapZebrain atlas. **(d)** Quantification of synaptic contact distribution in the anterior iIPN. Top: Distribution of single axons. Each row represents a dHb axon, and each column represents a bin along the mediolateral axis. Color coding represents relative frequency (from light to dark orange). dHb axons are sorted by the total number of detected contacts. Bottom: Pooled frequency distributions. Each gray line is a single axon, and the mean is shown in black. **(e)** dPPR^contra^ (green) and dPPR^ipsi^ (cyan) along with iIPN (gray) neurons. **(f)** Top left: schematic of targeted photoactivations. Top right: Fraction of gad1b positive dPPR^ipsi^ (cyan bar) and dPPR^contra^ (green bar) neurons. Bottom: example photoactivation experiment.

Using a DBSCAN clustering algorithm based on dendrite centroid positions (*ε* =4.5 µm, min. samples=5 neurons), we identified three glomeruli comprising 7, 7, and 16 iIPN neurons, respectively, along with 12 outliers (Figure 3a second, third and fourth columns). Coregistration of the EM and imaging data highlighted the third cluster (blue neurons in Figure 3a) as the anatomical substrate of our previous imaging results (Figure 3b).

Then, we detected axo-dendritic contacts from 4 glomerular neurons and reconstructed their presynaptic partners and other neurons terminating in the glomerulus (see Methods). A significant portion of these inputs originated from the dHb (n=39 neurons; 35 from r-dHb and 4 from l-dHb, including fully and partially reconstructed neurons, Figure 3c). These axons wrap around the iIPN neuropil and innervate multiple subregions, including the area corresponding to the contralateral (not reconstructed) glomerulus. To quantify this observation, we identified synapses formed by the reconstructed axons within the anterior iIPN (Figure S8b; see Methods), including both the left and right glomerulus. We then plotted their distribution along the mediolateral axis (Figure 3d), confirming that dHb axons innervate both regions.

A second group of presynaptic partners consisted of unipolar neurons from the ipsilateral (n = 1 neuron) as well as contralateral (n = 5 neurons) dorsal prepontine region (dPPR^ipsi^ and dPPR^contra^), located in the dorsal anterior part of the hindbrain. Unlike dHb neurons, their axon terminals had tufted endings resembling iIPN dendrites and exclusively terminated on the seed glomerulus (Figure 3e and Figure S8c). While these two populations where similar in morphology, we detected axo-dendritic contacts with iIPN neurons only for the dPPR^contra^ population.

To validate our EM findings, we performed targeted photoactivation experiments (Figure S9a) using animals coexpressing a photoactivable GFP variant (paGFP) and red-labeled dHb axons (n = 4 animals)^27^. We used the red channel to locate the glomerulus and activated the paGFP at 800 nm to retrogradely or anterogradely label neurons with axons or dendrites in the activation region, respectively. Consistent with EM reconstructions, we labeled the somata and axons of iIPN neurons (Figure S9b, panels I-III), dHb axons (Figure S8b, panel I), dPPR^ipsi^ and dPPR^contra^ neurons (Figure S9c, Figure S9b panels IV and V). While the somata of dPPR^contra^ neurons were confined to a smaller region, we detected a similar number of neurons in the two prepontine population (Figure S9c). We then repeated the photoactivation experiments in fish coexpressing paGFP and red-labeled GABAergic (gad1b promoter, n = 4 animals, Figure S8e) or glutamatergic neurons (vglut2a promoter, n = 3 animals, Figure S8f) and found that while dPPR^contra^ show a poor overlap for both markers, dPPR^ipsi^ are predominantly GABAergic (75% overlap with the gad1b marker, Figure 3g and Figure S8g).

To summarize, we found that glomerular iIPN neurons receive three distinct inputs: dHb (predominantly from the temperature-responding right subnucleus), dPPR^ipsi^ and dPPR^contra^ (Figure S8d). The dHb is known to be a glutamatergic region and is responsible for the thermal reward excitation component observed in downstream iIPN neurons^28^. In contrast, dPPR^ipsi^ neurons transmits GABAergic signals in the glomerulus, the role of which is still unclear (Figure 3h).

### Compartmentalized activity in dHb axon terminals mirrors iIPN glomerular tuning

We next imaged dHb axon terminals at the same location where dendritic iIPN glomerular activity had previously been recorded (Figure 4a, n = 9 animals). Based on dHb axon morphology and somatic activity patterns in dHb neurons, we expected symmetrical increases in dHb axon activity within the regions corresponding to both glomeruli during temperature stimulation. However, this is not what we found.

**Figure 4.**
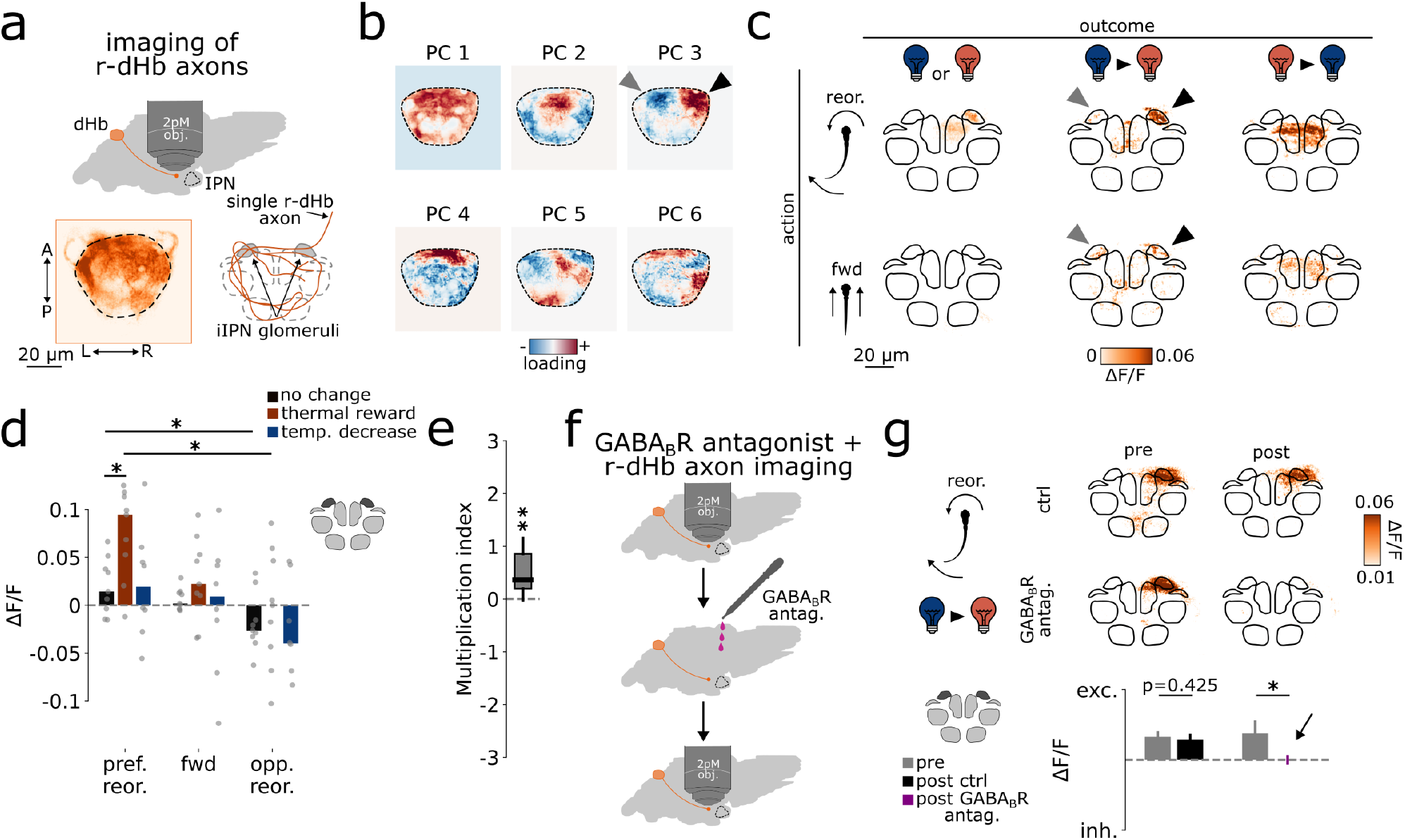
Integration of actions and outcomes arises in dHb axon terminals and is supported by presynaptic GABA_B_ receptors. **(a)** Top: Schematic of 2-photon calcium imaging experiments in dHb axons. Left bottom: Sum projection from an example recording. Right bottom: Schematic of dHb axon morphology. **(b)** Pixel-wise PC analysis. Loadings of the first 6 PCs are shown from blue (negative) to red (positive). **(c)** Average ΔF/F projection across action-outcome combinations. Rows represent actions, and columns represent outcomes. CCW and CW swims have been pooled together by flipping the stack along the midline. In this way, responses for the preferred reorientation can be seen on the right side, while responses to the opposite one are on the left. **(d)** Anatomical-based quantification. Blue: temperature decrease. Brown: thermal reward. Black: no temperature change. pref. reor.: contralateral reorientation relative to the underlying glomerulus’ anatomical position. fwd: forward (fwd) swim. opp. reor.: ipsilateral reorientation relative to the underlying glomerulus’ anatomical position. Each dot represents a single fish, with bars indicating the median across fish. **(e)** MultiplicationindexfordHbaxonpatches. **(f)** SchematicofpharmacologyexperimentsindHbaxons. **(g)** dHbaxon activity upon GABA_B_R antagonist treatment. Top: Average ΔF/F projection during reorientation followed by thermal reward. Rows represent experimental groups (first row: ctrl group, second row: treatment group) and columns represent experimental timepoint (first column: first imaging session, second column: second imaging session). Bottom: ΔF/F quantification (mean ± s.e.m.) for different experimental groups. (gray bars: first imaging session, black bar: second imaging session ctrl group, purples bars: second imaging session treatment group).

Population recordings revealed heterogeneous activity with marked lateralization^29^. Pixel-wise principal component analysis (PCA, see Methods) highlighted this heterogeneity, showing “patchy” components (Figure 4b). Some components were symmetric around the midline (PC1, PC2, PC4), whereas others were strongly asymmetric (PC3, PC5), with the patchy regions overlapping the underlying glomeruli. Analysis of the pixel-wise ΔF/F responses for different action-outcome combinations revealed a pattern similar to that observed in the glomeruli (Figure 4c, gray and black arrows in the second column). We then extracted the activity from the glomeruli-like regions and found the same anatomical action tuning (Figure 4d) and a similar multiplication index (Figure 4e) to that obtained in iIPN neurons. Interestingly, we also observed a second medial patch responding to temperature decrease (Figure 4c, light and dark blue arrows in the third column). This activity is likely related to a second pair of glomeruli with opposite outcome tuning with respect to the one studied here.

The observed activity footprints may originate for two reasons. The first is due to a small population of dHb neurons that encode action-outcome associations and relay this information to the glomerulus. However, this scenario is unlikely as we did not observe this tuning in dHb somatic activity and none of our reconstructed dHb axons project to a specific glomerulus, a result also consistent with previous studies^25,30^. Furthermore, similar patchy structures have been observed in the dorsal and ventral IPN^25,31^. The second and more plausible explanation, reconciling functional and anatomical data, is that dHb axon terminals are locally modulated^32^.

### GABA_B_ receptors modulate dHb axon terminals

A simple model that could account for the multiplicative activity observed in dHb axons proposes that reward information is transmitted by (r-)dHb neurons and this activity is locally modulated at the axon terminals by a hypothetical population of action-tuned neurons, driven by the preceding movement. This enhanced axonal activity, in turn, drives iIPN firing (Figure S10a).

One potent modulator of habenular glutamate release, identified in both zebrafish^32^ and mammals^33–35^, is the presynaptic GABA_B_ receptor (GABA_B_R). Interestingly, in mammals, GABA_B_R activation does not suppress neurotransmitter release^36^; rather, it strongly enhances it, leading to multiplicative-like responses at the axon terminals for the same level of somatic depolarization in habenular neurons^33–35^.

We tested the role of GABA_B_R first using a receptor antagonist (CGP 46381; see Methods). Fish underwent an initial imaging session, after which we applied the antagonist (treatment group, n = 7) or water (control group, n = 9) and re-imaged dHb axons (Figure 4f). The treatment had no effect on swim frequency (Figure S10b) nor global dHb axon activity, as assessed by pixel-wise ΔF/F standard deviation over the entire experiment (see Methods, Figure S10c-d). We then quantified its effect on the reward-tuned patch. Strikingly, GABA_B_R blockade abolished action-outcome responses (Figure 4g). This effect cannot be attributed to a general reduction in sensory responses in r-dHb somata since the percentage and response magnitude of thermal reward-tuned neurons assessed in the r-dHb subnucleus remained unchanged after treatment (n = 6 per group, Figure S11d-f), indicating that the impairment occurs at the level of the axon terminals.

We then tested the opposite manipulation by activating GABA_B_R using the agonist baclofen (Figure S11g, see Methods). To isolate the effect of GABA_B_R from other movement-related neuromodulation, we presented temperature stimuli in open-loop. As expected from iIPN data, open-loop responses were overall weaker and more variable than in the previous closed-loop condition. Nevertheless, agonist enhanced the peak responses in the two reward-tuned patches (Figure S11g).

Together, these findings suggest an important role for GABA_B_Rs in modulating r-dHb axon terminals and, consequently, in enabling action-outcome associations in iIPN neurons.

### GABAergic prepontine neurons convey action signals

Our previous findings strongly suggest that GABAergic transmission in the glomerulus modulates dHb axon activity according to the animal’s movements. Given that both dHb axons and iIPN neurons are glutamatergic, we imaged GABAergic neuropil within the glomeruli, corresponding to dPPR^ipsi^ axon terminals (Figure 5a-b, n = 13 animals). Consistent with our prediction, we found that these axons responded to directional swims (Figure 5c). While we still observed weak temperature responses, the interaction between sensory and motor variables was not multiplicative and the motor component dominated (Figure 5d-e). Action tuning mirrored that of iIPN neurons, with excitation for the contralateral direction relative to the glomerular hemisphere (CCW for the right dPPR^ipsi^ axons, CW for the contralateral counterpart) and inhibition for the opposite reorientation. The observed inhibition from baseline during non-preferred reorientations suggests that these neurons have a non-zero baseline firing rate. This, in turn, implies that dPPR^ipsi^ may modulate dHb axon terminal activity in a tonic manner, enhancing sensory transmission following preferred choices and weakening it after non-preferred ones.

**Figure 5.**
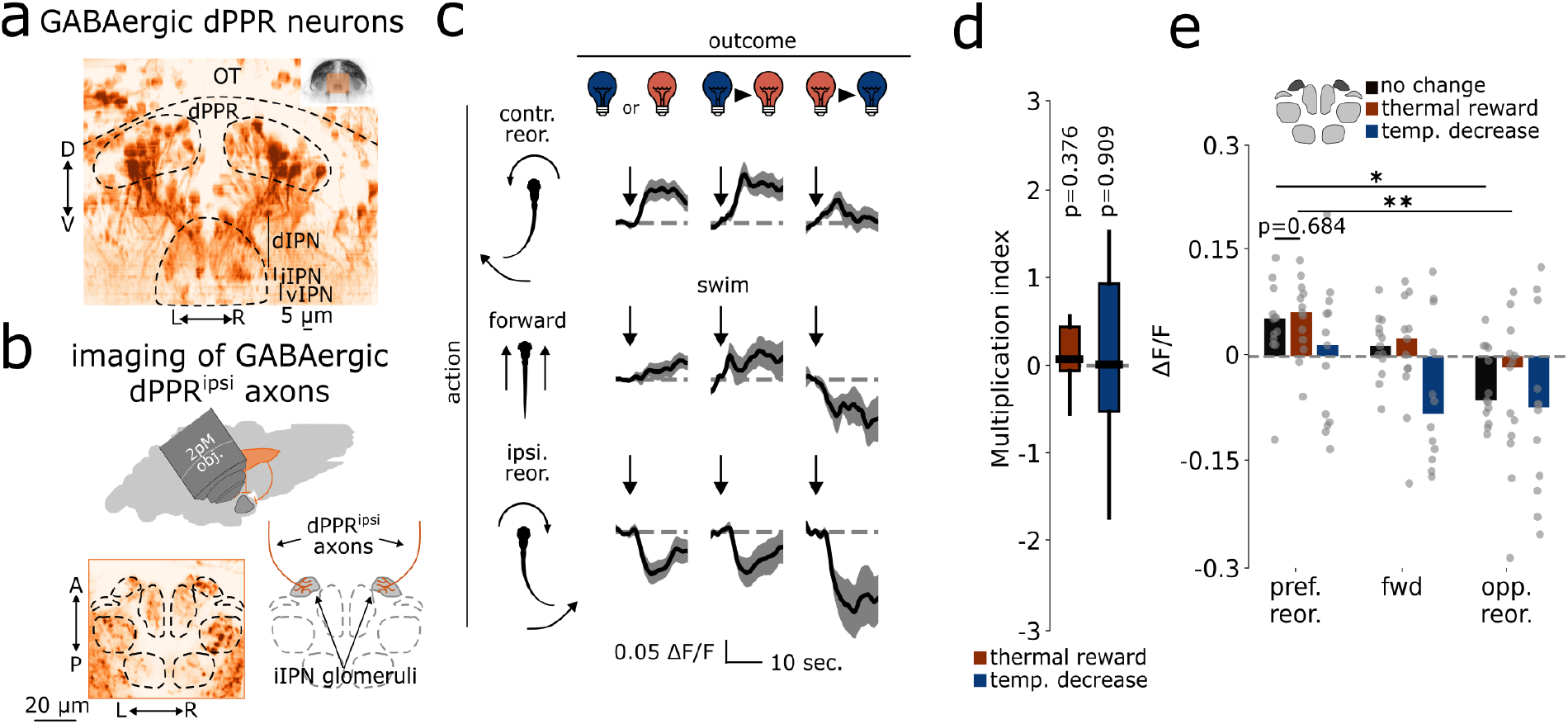
GABAergic prepontine neurons convey action signals to the iIPN glomeruli. **(a)** Frontal projection of GABAergic neurons targeting the IPN. **(a)** Top: Schematic of 2-photon calcium imaging experiments of dPPR^ipsi^ GABAergic axons. Bottom left: Sum projection from an example recording. Bottom right: Schematic of dPPR^ipsi^ projections. **(c)** Triggered average for different action-outcome combinations (mean ± s.e.m.). **(d)** Multiplication index when considering either temperature increase (brown) or decrease (blue). **(e) Anatomical**-based quantification. Blue: thermal reward. Brown: temperature decrease. Black: no temperature change. pref. reor.: contralateral reorientation relative to the underlying glomerulus. fwd: Forward swim. opp. reor.: ipsilateral reorientation relative to the underlying glomerulus position. Each dot represents a single fish, with bars indicating the median across fish.

## Discussion

In the present study, we describe how the brain of larval zebrafish links actions to a behaviorally relevant sensory outcome.

Using an operant thermoregulatory assay^8–14^, we found that fish create short-term associations between actions and thermal reward to avoid cold temperatures. Associations were completely abolished when we ablated the dHb, suggesting that the dHb-IPN pathway is crucial in establishing sensorimotor associations between the animal and its environment. These results are in line with previous studies linking this pathway to fear and avoidance learning^18–21,33,37,38^, experience-dependent navigation^16^, learning of directional rules^39^ and generally support the idea of the dHb as an experience-dependent switchboard of behavior^7^.

We next found that sensory and motor information are integrated in the IPN through the convergence of the dHb^25,40,41^ and dPPR pathway^29,40,42–47^(Figure 6a). Interestingly, the integration time window observed in iIPN neurons aligns with that identified in the operant assay through the delay experiments. One interpretation could be that these iIPN neurons underlie the establishment of an internal representation of the last action-outcome relationship (i.e. a memory trace, Figure 6b). This representation would then allow the animal to, through a yet undetermined mechanism, bias turning behavior in response to future aversive temperature changes. This idea is consistent with a previous study linking the region to evidence accumulation to favor swims of particular direction. More broadly, IPN neurons may integrate sensory input, internal state, sensorimotor contingencies and perhaps other cognitive variables to adaptively guide turning behavior^29,31,48,49^.

**Figure 6.**
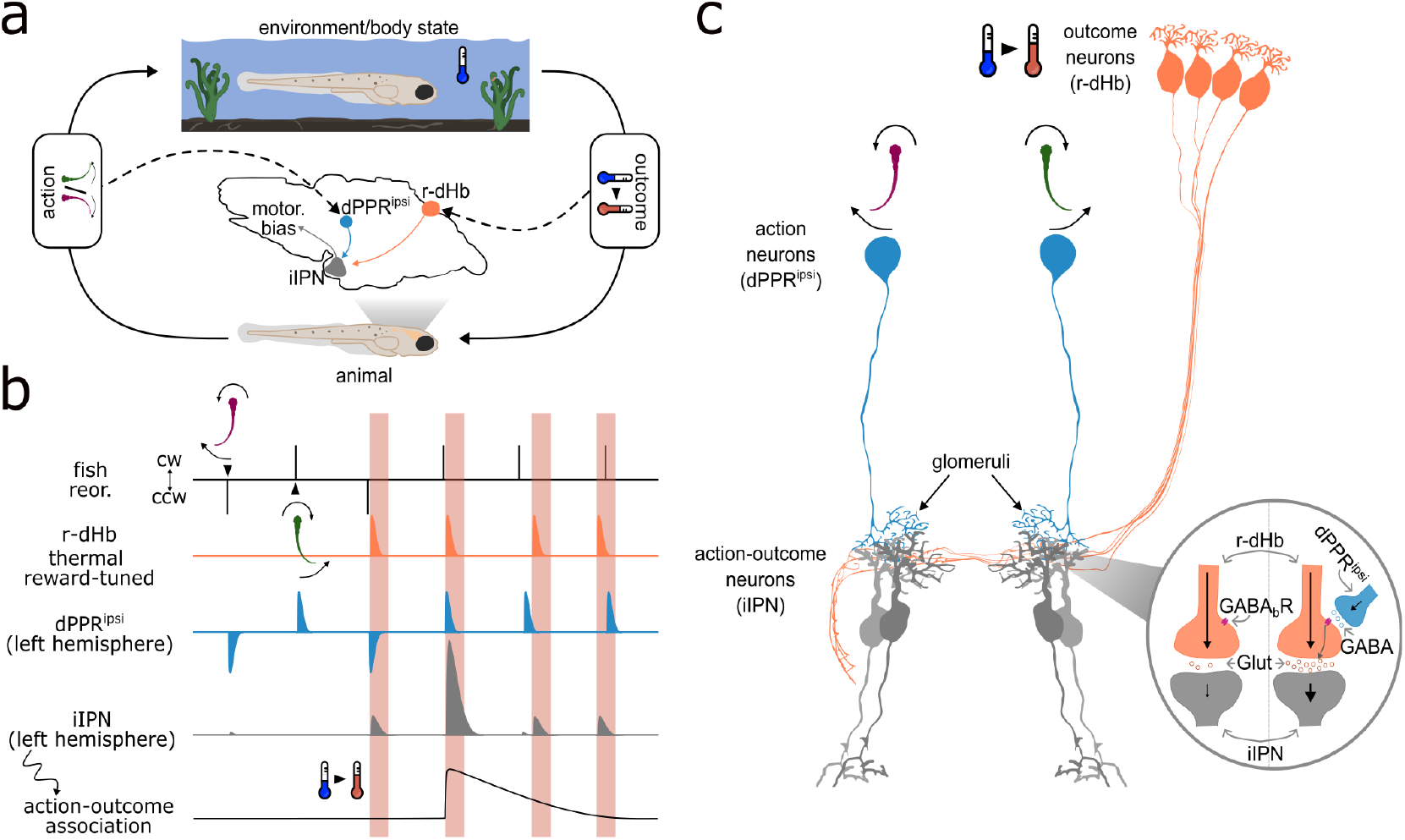
Summary.

One important question for future research is to assess whether these neurons are specialized for integrating temperature changes or are instead multimodal. The latter hypothesis seems more likely, as it is known that dHb projections carry odor information^50–53^ and other sensory variables^16,54–56^. Another question is how neurons downstream of the iIPN bias behavior. Considering the strong connections between this region and some monoaminergic systems, an hypothesis could be that neurotransmitters such as dopamine and serotonin could be involved^57–59^.

Strikingly, we found that the multiplication of actions and thermal reward does not arise in iIPN neurons but rather in the dHb presynaptic axon terminals (Figure 6c). This mechanism offers an elegant and efficient solution for linking actions and outcomes, requiring only one sensory neuron, with different axon segments modulated independently by distinct animal’s decisions. For instance, here a single dHb axon could target both the left and right iIPN glomerulus through en passant synapses and its segments selectively modulated by different movements. A similar mechanism has been described in the insect mushroom body, where dopaminergic neurons modulate segments of olfactory axons in a behavioral context-dependent manner^60^.

Finally, we found that this multiplication is supported by GABA_B_R, acting as positive gain modulator of dHb signals in response to animal’s movements. Our results are in line with mouse studies showing that this receptor enhance glutamate release from the medial habenula (mHb, the homolog of the zebrafish dHb)^33,34,61–63^. In mouse, this enhancement is both IPN subregion-^64^ and cell-type-specific^35^. A similar process appears to occur in larval zebrafish, where recent findings have shown that GABA_B_R signaling in the dIPN reduces dHb-to-IPN transmission, whereas in the iIPN it appears to enhance it^32^. Importantly, this does not exclude additional mechanisms, such as post-synaptic inhibition^59^, intrinsic properties of iIPN neurons, retrograde inhibition^65^ or other forms of neuromodulation^66,67^.

Even though the mammalian brain have far more neural resources than the larval zebrafish one, the mechanism identified here may serve a similar function, a hypothesis that can be experimentally tested. To conclude, while some implementation details may be unique to the dHb-IPN pathway, the modulation of reward-related axons by choice-selective neurons may represent a general principle employed by other brain circuits performing similar tasks^2–5,68–72^.

## Acknowledgements

The authors thank the members of the Portugues lab for constructive discussions throughout the project. In particular, we want to thank You Kure Wu for the constant technical and scientific support. We also thank Thomas Frank, Thomas Misgeld and especially Ilona Grunwald Kadow for the support and constructive feedback received during the project. This research was funded by the German Research Foundation (DFG) through SPP 2205 (project 430156228), Germany’s Excellence Strategy within the framework of the Munich Cluster for Systems Neurology (EXC 2145 SyNergy, identifier 390857198), through the “Enhanced resolution microscopy” (project 518284373), by the Volkswagen Stiftung via a Life? grant and by the Max Planck Gesellschaft.

## Contributions

EP, VP and RP designed the project. EP built the IR laser freely swimming and head-fixed rigs with support from VP. EP performed the experiments, with the support of VP (during iIPN somata imaging and initial freely-swimming experiments) and the support of AS (for dHb chemogenetic ablations and anatomies). EP performed all analysis and quantifications. EM reconstructions were done by ariadne.ai AG and in a small amount by EP. EP, VP and RP wrote the manuscript.

## Materials and Methods

### Zebrafish husbandry

All procedures related to animal handling were conducted following protocols approved by the Technische Universität München and the Regierung von Oberbayern (TVA 55-2-1-54-2532-101-12 and TVA 55.2-2532.Vet_02-24-5). Adult zebrafish (Danio rerio) from Tüpfel long fin (TL) strain were kept at 27,5-28°C on a 14/10 light cycle, and hosted in a fish facility that provided full recirculation of water with carbon-, bio-and UV filtering and a daily exchange of 12% of water. Water pH was kept at 7,0-7,5 with a 20 g/liter buffer and conductivity maintained at 750-800 μS using 100g/liter. Fish were hosted in 3,5 liter tanks in groups of 7-17 animals and fed with Gemma micron 300 (Skretting USA) and live food (Artemia salina) twice per day. Larvae were fed with Sera micron Nature (Sera) and ST-1 (Aquaschwarz) three times a day. All experiments were conducted on 5-8 dpf larvae of yet undetermined sex. The week before the experiment, one male and one female or three male and three female animals were left breeding overnight in a sloping breeding tank or breeding tank (Tecniplast). The day after, eggs were collected in the morning, rinsed with water from the facility water system, and then kept in groups of 20-40 in 90 cm Petri dishes filled with Danieau solution 0.3x (17.4 mM NaCl, 0.21 mM KCl, 0.12 mM MgSO_4_, 0.18 mM Ca(NO_3_)_2_, 1.5 mM HEPES, reagents from Sigma-Aldrich) until hatching and in water from the fish facility afterwards. Larvae were kept in an incubator at 28,5°C and a 14/10 hour light/dark cycle, and their solution was changed daily. At 4 or 5 dpf, animals were lightly anesthetized with Tricaine 440 mesylate (Sigma-Aldrich) and screened for fluorescence under an epifluorescent microscope.

### Animal strains

The Tuepfel long-fin (TL) wild-type strain was used for most of freely swimming experiments. For imaging experiments, fish were mitfa -/- (i.e., nacre) mutants lacking melanophores to allow optical access to the brain. Transgenic lines used are:

- Tg(16715:Gal4);Tg(UAS:Ntr-mcherry)^16^ used for dHb chemogenetic ablation and targeted photoactivation experiments
- Tg(16715:Gal4);Tg(UAS:GCaMP6s)mpn101^16,73^ used for dHb somata and axons imaging experiments and anatomies
- Tg(elavl3:GCaMP6s)^74^ used for dHb somata imaging experiment
- Tg(evavl3:Hsa.H2B-GCaMP6s)^75^ used for iIPN and dPPR somata imaging experiments
- Tg(s1168t:Gal4);Tg(UAS:GCaMP6s)mpn101^73,76^ used for iIPN dendrites (glomeruli^iIPN^) imaging experiment and anatomies
- Tg(gad1b:Gal4);Tg(UAS:GCaMP6s)mpn101^73,77^ used for dPPR axon (glomeruli^dPPR^) imaging experiments and anatomies
- Tg(Cau.Tuba1:c3paGFP)a7437^27^ used for photoactivation experiments
- Tg(gad1b:RFP)^78^ for anatomy and targeted photoactivation experiments.
- Tg(vglut2a:DsRed)^79^ for anatomy and targeted photoactivation experiments.

### Experimental Setups

#### IR laser freely swim setup

This setup was inspired by a previous publication^17^ and largely follow the same design. Briefly, fish were placed in an arena 200 mm x 110 mm (preference assay) or 200 mm x 65 mm (operant thermoregulatory assay), illuminated with 750 nm IR illumination, and allowed to move freely. For all experiments, we projected a visual pattern (Optoma) on the arena floor, following pink noise statistics in spatial frequencies, to provide the fish with additional self-motion cues. Fish were tracked at 150 Hz (camera: ximea USB3.0 MQ022RG-CM, objective: 25 mm Edmund Optics objective) using Stytra^16,29,80^, from which body position and heading were extracted. Positional data controlled the angles of two scan mirrors (Thorlabs, GVS102) using a DAQ board (National Instruments, 781438-01), maintaining a 980 nm laser beam centered on the fish’s center of mass. Based on the experimental conditions, the laser beam was either centered on the fish (thermal reward condition) or redirected to a distant heat sink outside the behavioral arena. This approach minimized delays and facilitated sharp temperature changes. In front of the camera we additionally placed a 800 nm shortpass filter (Thorlabs, ESH0800) and a 650 nm longpass filter. For calibrations, we fixed a thermocouple^16^ to the behavioral arena and filled it with water. Then, the laser beam was rapidly moved on the thermocouple or on a 5 mm distant region. Temperature was recorded with a temperature controller (Warner Instruments, Cl 100) connected to the behavioral PC through a LabJack series U3-LV (LabJack)^16^.

#### IR laser setup for two-photon imaging

The same IR laser was positioned in front of a 1-axes galvanometric mirror, which switched between two states. In one state, the mirror directed the beam toward the fish, positioned frontally at an angle that avoided the objective and allowed the beam to reach the fish body. In the second state, the mirror redirected the beam to a heat sink distant from the animal. To filter out potential contaminants detected by the microscope’s photomultipliers, we placed a 750 nm long-pass filter (Thorlabs, FELH0750) in front of the IR laser. In order to avoid temperature accumulations, the laser power used was reduced to 1/3 of the one used for freely swim experiments. Furthermore, the fish was perfused with 20 °C water as previously done^16^. We presented optic flow visual stimulation at 40 Hz (Asus P2E microprojector). A red long-pass filter (Kodak Wratten No.25) allowed the simultaneous 905 nm imaging and visual stimulation. Fish were illuminated using infrared light-emitting diodes (850 nm wavelength) and imaged from below at 200 Hz (Pike F032B, Allied Vision Technologies).

#### Two-photon microscopy

Functional imaging data were acquired at 2-5 Hz, depending on the calcium sensor kinetic being used in the experiment. Laser power and field of view were adjusted based on the particular structure being imaged. For anatomical data, we acquired 10/45 frames (1Hz rate for 10/45 seconds) per plane with a 1-2 microns inter-plane distance. Once a plane was imaged, the objective was moved by 1 micron and the acquisition restarted. Dual channel (red and green) anatomical stacks were acquired at 950-1000 nm while single channel ones (only green) at 905 nm.

### Animal preparation and behavioral protocols

All experiments where behavior was recorded were run from 10.00 h to 20.00 h, room temperature was set to 20/21 °C. All visual and thermal stimulus presentation as well as behavioral tracking were performed using the Stytra package^80^.

#### Cold avoidance experiment

Here, the IR laser targeting the fish was contingent on the fish being inside an experimentally defined rectangular region (ROI)^81^. Such a region was invisible to the fish and once inside, a temperature increase was produced through the IR laser.

#### Operant thermoregulatory assay

This protocol^11^ was designed to test whether zebrafish can associate a directional choice with a specific temperature consequence. The first experiment presented comprised 60 trials: 30 trials where an increase in temperature was paired with a turn in a designated ‘rule direction’ (either CCW or CW), and 30 trials in the opposite direction. For delay and dHb ablation experiments we did 10 trials per block, asprevious resultshighlighted a short-and not a long-term memory process building over many trials. When inside an experimentally defined region (not close to arena borders) a trial starts by decreasing the temperature (temperature decrease) at the end of a swim of any kind (CCW, forward or CW). Then, the animal can increase the temperature again by turning toward the rule direction. The trial ends when the fish turns in the rule direction or after 10 seconds from temperature decrease (timeout). Then, after a minimum ITI of 5 seconds (where the laser is directed toward the animal) and the fish being away from the border, the next trial starts. As this was a self-paced experiment, its duration varied between fish. All temperature transitions were triggered by the animal’s movement, as we observed a net decrease in swim rate otherwise^82^. Fish were tested once, between 5 and 8 days post-fertilization (dpf), and otherwise kept in an incubator at 28 °C near the behavioral apparatus. Performance did not significantly differ with age.

#### 2-photon imaging behavioral protocol

This protocol was designed to assess the combined effects of actions and thermal consequences on different neuronal populations. For imaging of iIPN somata, temperature changes were stochastically triggered at the end of swim events, with a 25% probability that each swim would switch the current laser state. In the rest of closed-loop imaging experiments, a similar protocol was applied, but once the laser turned ON, it remained ON for a minimum of 10 seconds. The first swim after this interval turned the laser OFF, initiating another 10 second long refractory period. After this, each swim had a 30% probability of turning the laser ON again. This modification ensured relatively frequent temperature increases while allowing sufficient time between changes to facilitate clear baseline extraction and quantification. Throughout the entire experiment we also presented whole field forward motion at 3 mm/s to promote swimming events with rotational as well as translational congruent visual feedback. Translational and rotational feedback were computed online as previously done^48^. The experiment lasted 30-60 minutes and only one plane was acquired per each fish. If during the experiment the acquisition drifted, the experiment was restarted for a maximum of three times.

#### Head-fixed preparation

For functional imaging experiments, fish were embedded in 2.5% low-melting point agarose in 30 mm petri dishes. The agarose around the tail and nose was removed with a scalpel to allow tail movements and direct stimulation of the olfactory bulb^16,55^. Once the animal was placed under the objective, we waited 5-10 minutes with the fish engaged in the same visual closed-loop that will experience later. For anatomical experiments, fish were embedded in 1.5%/2% low-melting point agarose in 30 mm petri dishes.

### Manipulations

#### Chemogenetic ablation

Targeted dHb ablation was performed using the Nitroreductase (Ntr)-Nifurpirinol (NfP) chemogenetic approach^83^. Ntr catalyzes the reduction of the pro-drug NfP into a cytotoxic compound, leading to cell death. By expressing Ntr in specific cells, targeted ablations can be achieved. We used the Tg(Gal4:16715):Tg(UAS:Ntr-mCherry) line^16^, which selectively expresses Ntr-mCherry in the dorsal habenula of larval zebrafish. Larvae were first screened for mCherry expression using an upright epifluorescence microscope. mCherry-positive larvae (ablation group) and mCherry-negative larvae (treatment control group) were then placed in separate 90 mm Petri dishes containing minimal water. Subsequently, 30 mL of a 5 µM NfP solution with 0.2% DMSO (diluted 500 times from a 2.5 mM NfP stock) was added to each dish. The Petri dishes were incubated in a dark box at 28°C for 16 hours to prevent degradation of the photosensitive NfP. Both the preparation of the NfP solution and the transfer of fish were also conducted in the dark to minimize NfP degradation. After the incubation period, larvae were removed from the solution and thoroughly washed three times to eliminate any residual NfP and DMSO. The fish were then left to recover in fish water for one day. To evaluate the efficiency of dHb ablation, random larvae were selected before and after treatment and imaged using a confocal microscope to assess mCherry expression in dHb neurons.

#### Pharmacology experiments

We used either the GABA_B_R antagonist CGP 46381 (50 µM, CAS 123691-14-5, Santa Cruz Biotechnology) or the GABA_B_R agonist (R)-Baclofen (150 µM, CAS 69308-37-8, Tocris). After a first imaging session (as described elsewhere), head-fixed fish were brought under a stereomicroscope close to the experimental rig. Then, we carefully washed for three times with either water (ctrl group) or agonist/antagonist (treatment group) using a pasteur pipette. Fish was subsequently moved again in the two-photon microscope rig, let habituate to the visual closed-loop and allow drug diffusion for 10 minutes, and started the second imaging session. In this experiment, bath application was necessary, as intraventricular injections completely abolish fish swimming behavior, particularly immediately after injection.

### Anatomical registration

Registrations were done using a combination of manual and automatic procedures using the free Computational Morphometry Toolkit (CMTK - http://www.nitrc.org/projects/cmtk) (volumetric data) or the python package dipy. Each single-plane dataset with the exception of r-dHb somata imaging have been morphed with an internal reference, to allow for average ΔF/F projections and quantifications. All volumetric data (EM, photoactivations, anatomies and multiplane dPPR population imaging) have been morphed to the Max Planck Zebrafish Brain Atlas mapzbrain (https://mapzebrain.org)^84^ for visualizations and quantifications.

### Data analysis

All analyses were performed using Python 3.8 and relevant Python libraries for scientific computing, like numpy, scipy and scikit-learn.

#### Preprocessing of behavioral data

Relevant behavioral parameters were tracked online using Stytra^80^. All the behavioral preprocessing was performed using the python package Bouter (https://github.com/portugueslab/bouter) and as previously described^16,29,48^. For head-fixed experiments reorientation angle was computed as the cumulative tail bending over the first 70 ms from swim onset^85^ while for freely swimming experiment the cumulative reorientation from swim start to end. To categorize CCW, forward and CW swims in head-fixed experiments we set a threshold of 10 degrees.

#### ROI occupancy

In Figure S1f and h, we report ROI occupancy as a measure of cold avoidance behavior. If the fish exhibit cold avoidance, they will spend more time in the warmed region. ROI occupancy was computed as the fraction of time spent inside the ROI over the entire experiment duration, multiplied by 100.

#### Outside ROI exposure time

In Figure S1g, we report the outside ROI exposure time as another measure of cold avoidance. For each instance, we computed the time spent from ROI exit to re-entry and then averaged this value per fish.

#### Angular velocity

In Figure S2c, we report the angular velocity at temperature transitions during the operant assay. Angular velocity was computed by first smoothing the fish heading using a Savitzky–Golay filter (window = 150 ms), then calculating the first-order derivative and dividing it by the time interval between consecutive timepoints. Conditioned responses were evaluated by measuring the average angular velocity (baseline-subtracted) from 0.5 to 0.9 sec. after temperature decrease.

#### Reorientation bias

In Figure 1c,f,i and Figure S2e,f we report reorientation bias as an index of task performance. This was defined as the average reorientation angle of the first swim detected after temperature decrease for trials where the preceding ITI was lower (or higher for Figure S2f, yellow) than 7.5 sec (except for Figure 1d). For Figure S2e we considered all the swims during ITIs. For shuffle controls, we shuffled rule block’s identity.

#### Reorientation magnitude

In Figure S3b we report reorientation magnitude as an index of temperature-induced innate turning behavior. This was defined as the average absolute reorientation angle of the first swim detected after temperature decrease for trials where the preceding ITI was lower than 7.5 sec. For shuffle controls, we randomly selected swims throughout the experiment.

#### Cold exposure time

In Figure 1e,h we report cold exposure time as a measure of task performance. This was defined as the average time spent from temperature decrease to thermal reward for trials where the preceding ITI was lower than 7.5 sec. For delay experiments, this value was corrected by the applied delay to represent cold exposure time under a zero-delay condition.

#### Preprocessing of imaging data

All functional data were drift-corrected using the Python package fimpy (https://github.com/portugueslab/fimpy) or suite2p^86^. ROIs were manually extracted using napari (https://napari.org/stable/) based on clear anatomical landmarks, except for habenula somata (extracted using suite2p). Each ROI was resampled at 5 Hz with linearly spaced timepoints. Baseline fluorescence (F0) was defined as the median over the entire experiment. ΔF/F was computed as (F-F0)/F0, where F is the instantaneous fluorescence from the raw ROI time series. These time series were then smoothed with a third-order Savitzky–Golay filter over 11 frames.

#### Fraction of temperature-responding r-dHb neurons

In Figure S5d and Figure 4i, we report the fraction of temperature-modulated r-dHb neurons. Only positive modulations (increases from baseline fluorescence) were considered. To determine this fraction, we first calculated the evoked using a 10-second window centered on stimulus onset or offset (each condition analyzed separately). Baseline activity was subtracted, and the average response was calculated. This procedure was repeated 1000 times, each time circularly shuffling the original trace to create a null distribution of evoked activity. A p-value was computed per neuron, and the fraction of responsive neurons was defined as the proportion with p-values below 0.05.

#### r-dHb thermal reward response dynamics

Related to Figure S5e-g r-dHb thermal reward response dynamics were classified by first computing the response triggered average. We then extracted the ΔF/F from 0–2 sec. and 8–10 sec. These values were used to calculate a polar angle describing the relative ΔF/F over the two time windows. For example, a fast-adapting neuron with a greater increase in the first window compared to the second would have a polar angle close to zero, whereas a slow-adapting neuron would have an angle close to 90 degrees. Arbitrary thresholds were then set to categorize response profiles.

#### Multiplication index

Related to Figure 2f, Figure 4e, Figure S5d and Figure S7b,c. For iIPN and r-dHb somata imaging, we computed this index only on the subset of neurons significantly modulated by thermal reward while for the rest of the experiments we directly computed it on the two traces extracted from the glomerular region (or corresponding dHb axon patches). We then computed the triggered average for every action-outcome combination and calculated the maximum ΔF/F increase over a 10-sec window. The multiplication index was then computed as the negative log-ratio between: (1) the difference in between CCW and CW swims in the absence of thermal feedback and (2) the difference in between CCW and CW swims when they led to thermal reward.

#### Anatomy-based choice-outcome responses

Related to Figure 2e, Figure 4d Figure 5e. For every neuron we computed the triggered average for every action-outcome combination and calculated the average ΔF/F increase over a 10 sec. window. Then, we pooled CCW (divided by outcome responses) responses for neurons located on the right brain hemisphere with CW responses for the ones located on the left hemisphere and did the same for the opposite combination. Then we computed the median across neurons within the same animal.

#### Sensorimotor lag

Related to Figure 2j. During open-loop stimulation, each thermal reward instance was assigned a sensorimotor lag value. We searched for the closest swim event within a 2-second window before (−2 sec) and after (+2 sec) thermal reward. Sensorimotor lag was defined as the difference between the timing of the first swim onset and thermal reward. Trials where the sensorimotor lag was positive but the fish swam at least 5 seconds before thermal reward were excluded. Finally, we did the same pooling criterion used for the “Anatomy-based choice-outcome responses”. Sensorimotor lag values were finally binned with 0.5-sec. spacing.

#### Analysis ΔF/F standard deviation

Related to Figure S9c-d. Starting from the ΔF/F aligned stacks (time,y,x), we computed, for every pixel, the standard deviation over time. In this way, we obtained a 2D map (y,x). For Figure S9c we then calculated the rigid transformation matrix (RigidTransform2D class from the python package dipy) for all the acquisitions using aligned stack sum projections as a common reference (a selected recording). Finally, we transformed all standard deviation maps and averaged across fish. For Figure S9d we then obtained, from each map, the average standard deviation value in dHb axons by averaging in space and calculated the ratio for first and second imaging session across conditions.

#### Pixel-wise PCA

Related to Figure 4b. Imaging experiment can be seen as a 3-dimensional matrix (t,y,x). We flattened this matrix to obtain a bi-dimensional matrix (t,y*x) and performed PCA along the y*x dimension. Then, from the PCA loadings we reconstructed the original anatomical image to generate a loading map for the first 6 PCs.

### Electron microscopy reconstructions

EM reconstructions were performed on an open-access Serial Blockface EM dataset from a 5 dpf larval Tg(elavl3:GCaMP5G)a4598 sampled at a resolution of 14 × 14 × 25 nm^26^. The IPN region was identified based on prior work^25,48^. Reconstructions were generated by merging neurite segments identified by an automated flood-filling algorithm as described in^26^. The first reconstructed neurons were from the iIPN neurons, traced by annotators at ariadne.ai AG, starting from seed points selected by the authors. Each completed cell underwent a quality review by an expert annotator. From the aggregated mesh, neuron skeletons were extracted using custom python software. Presynaptic partners were identified by first detecting synapses with a detection algorithm^26^, followed by manual curation of contact points (as done elsewhere^25,87^) and tracing of partner neurons. Partner reconstruction and quality control were performed by both ariadne.ai AG and the authors. Unlike iIPN, reconstructing partner neurons was more challenging due to neurites extending into blurred areas of the dataset, making this reconstruction effort incomplete. Similarly, many dHb axons are likely incomplete, due to a discontinuity in the dataset in the posterior-ventral-left IPN. Left- and right-dHb afferents were distinguished by the side of the fasciculus retroflexus along which each reconstructed axon traveled. Reconstructions classification was done manually based on dendritic and axonal positioning. Cellular compartments (glomeruli and axons) were identified by morphological characteristics such as process thickness. iIPN glomeruli were classified using the DBSCAN clustering algorithm from the sklearn library in Python.

### Statistics

- Figure 1c: reorientation bias CCW block: - 0.093 ± 0.037 rad., mean ± s.e.m. vs. CW block: + 0.130 ± 0.040 rad., mean ± s.e.m.; two-sided Wilcoxon signed-rank test, p<0.0001; CCW shuffle block: + 0.013 ± 0.033 rad., mean ± s.e.m. vs. CW shuffle block: + 0.121 ± 0.032 rad., mean ± s.e.m.; two-sided Wilcoxon signed-rank test, p=0.86; CCW block vs. CCW shuffle block, two-sided Wilcoxon signed-rank test, p<0.01; CW block vs. CCW shuffle block, two-sided Wilcoxon signed-rank test, p < 0.001
- Figure 1e: exposure time 0 sec. delay: + 1.985 ± 0.171 sec., mean ± s.e.m., exposure time 0.4 sec. delay: + 3.19 ± 0.388 sec., mean ± s.e.m., exposuretime 0.8 sec. delay: + 3.432 ± 0.418 sec., mean ± s.e.m.. 0 delay vs. 0.4 delay, two-sided T-test, p < 0.01, 0 delay vs. 0.8 delay, two-sided T-test, p < 0.01, 0.4 delay vs. 0.8 delay, two-sided T-test, p = 0.685
- Figure 1f: reorientation bias 0 delay: + 0.231 ± 0.070 rad., mean ± s.e.m., reorientation bias 0 delay shuffle: - 0.0005 ± 0.006 rad., mean ± s.e.m., reorientation bias 0.4 delay: + 0.106 ± 0.044 rad., mean ± s.e.m., reorientation bias 0.4 delay shuffle: + 0.01 ± 0.006 rad., mean ± s.e.m., reorientation bias 0.8 delay: - 0.053 ± 0.07 rad., mean ± s.e.m., reorientation bias 0.8 delay shuffle: + 0.004 ± 0.006 rad., mean ± s.e.m.. 0 delay vs. 0 delay shuffle, two-sided Wilcoxon signed-rank test, p < 0.001, 0.4 delay vs. 0.4 delay shuffle, two-sided Wilcoxon signed-rank test, p = 0.114, 0.8 delay vs. 0.8 delay shuffle, twosided Wilcoxon signed-rank test, p = 0.231
- Figure 1h: exposure time ctrl: + 2.355 ± 0.147 sec., mean ± s.e.m., exposure time ablation: + 3.069 ± 0.270 sec., mean ± s.e.m., exposure time 0.8 sec. delay: + 3.432 ± 0.418 sec., mean ± s.e.m.. ctrl vs. ablation, two-sided T-test, p < 0.05
- Figure 1i: reorientation bias ctrl: + 0.127 ± 0.05 rad., mean ± s.e.m., reorientation bias ctrl shuffle: + 0.002 ± 0.006 rad., mean ± s.e.m., reorientation bias ablation: + 0.030 ± 0.061 rad., mean ± s.e.m., reorientation bias ablation shuffle: + 0.030 ± 0.061 rad., mean ± s.e.m.. ctrl vs. ctrl shuffle, two-sided Wilcoxon signed-rank test, p<0.01, ablation vs. ablation shuffle, two-sided Wilcoxon signed-rank test, p = 0.397
- Figure 2e: pref. reor. + no change: 0.04, median, pref. reor. + thermal reward: 0.288, median, pref. reor. + temperature decrease: -0.051, median, fwd + no change: -0.04, median, fwd + thermal reward: 0.164, median, fwd + temperature decrease: -0.073, opp. reor. + no change: -0.009, median, opp. reor. + thermal reward: 0.099, opp. reor. + temperature decrease: -0.008, pref. reor. + no change vs. opp. reor. + no change, two-sided Wilcoxon signedrank test, p < 0.001, pref. reor. + thermal reward vs. opp. reor. + thermal reward, two-sided Wilcoxon signed-rank test, p < 0.001, pref. reor. + no change vs. pref. reor. + thermal reward, two-sided Wilcoxon signed-rank test, p < 0.001
- Figure 2f: MI glomerulus: 1.413, median, two-sided Wilcoxon signed-rank test, p < 0.001
- Figure 2h: ΔF/F temp. incr. ctrl: 0.44, median, ΔF/F temp. incr. ablation: 0.16, median; ctrl vs. ablation, two-sided Mann-Whitney U test, p < 0.001
- Figure 2j: ΔF/F glom. pref. reor. (from left to right): 0.034, 0.056, 0.079, 0.184, 0.0156, 0.0452, 0.0714, 0.063, mean. ΔF/F glom. opp. reor. (from left to right): 0.041, 0.08, 0.0816, 0.076, 0.033, 0.067, 0.093, 0.0761, mean. ΔF/F glom. pref. reor. [-0.5,0] vs, ΔF/F glom. pref. reor. [0,0.5], two-sided Mann-Whitney U test, p < 0.05
- Figure 2k: ΔF/F only motor: + 0.068 ± 0.013, mean ± s.e.m., ΔF/F only sensory: + 0.289 ± 0.048, mean ± s.e.m., ΔF/F sensory + motor (arithmetic sum): + 0.358 ± 0.053, mean ± s.e.m., ΔF/F sensory + motor (observed): + 0.563 ± 0.098, mean ± s.e.m.. ΔF/F sensory + motor (arithmetic sum) vs. ΔF/F sensory + motor (observed), two-sided Wilcoxon signed-rank test, p < 0.01
- Figure 3f: median dPPR_ipsi_: 0.78, median dPPR_contra_: 0.25.
- Figure 4d: pref. reor. + no change: 0.014, median, pref. reor. + thermal reward: 0.095, median, pref. reor. + temperature decrease: 0.019, median, fwd + no change: 0.002, median, fwd + thermal reward: 0.022, median, fwd + temperature decrease: 0.009, opp. reor. + no change: -0.026, median, opp. reor. + thermal reward: 0, opp. reor. + temperature decrease: -0.04, pref. reor. + no change vs. opp. reor. + no change, two-sided Wilcoxon signed-rank test, p < 0.05, pref. reor. + thermal reward vs. opp. reor. + thermal reward, two-sided Wilcoxon signed-rank test, p < 0.05, pref. reor. + no change vs. pref. reor. + thermal reward, two-sided Wilcoxon signed-rank test, p < 0.05.
- Figure 4e: MI dHb patch: 0.366, median, two-sided Wilcoxon signed-rank test, p < 0.01
- Figure 4g: dHb patch (pref. reor. + thermal reward): pre ctrl vs. post ctrl: two-sided Wilcoxon signed-rank test, p = 0.425. pre treatment vs. post treatment: two-sided Wilcoxon signed-rank test, p < 0.05.
- Figure 5d: MI dPPR axon terminal (thermal reward): 0.072, median, two-sided Wilcoxon signed-rank test, p = 0.376. MI glomerulus^dPPR^ (temperature decrease): 0.014, median, two-sided Wilcoxon signed-rank test, p = 0.909.
- Figure 5e: pref. reor. + no change: 0.054, median, pref. reor. + thermal reward: 0.063, median, pref. reor. + temperature decrease: 0.016, median, fwd + no change: 0.015, median, fwd + thermal reward: 0.025, median, fwd + temperature decrease: - 0.082, opp. reor. + no change: -0.063, median, opp. reor. + thermal reward: -0.015, opp. reor. + temperature decrease: -0.073, pref. reor. + no change vs. opp. reor. + no change, two-sided Wilcoxon signed-rank test, p < 0.05, pref. reor. + thermal reward vs. opp. reor. + thermal reward, two-sided Wilcoxon signed-rank test, p < 0.01, pref. reor. + no change vs. pref. reor. + thermal reward, two-sided Wilcoxon signed-rank test, p = 0.684.
- Figure S1f: ROI occupancy: + 67.593 %, median vs. ctrl: + 24.321 %, median; two-sided Mann-Whitney U test, p < 0.01
- Figure S1g: outside ROI exposure time: + 0.68 sec., median vs. ctrl: + 32.325 sec., median, two-sided Mann-Whitney U test, p < 0.05
- Figure S1h: ROI occupancy first half: + 68.205 %, median vs. second half: + 67.515 %, median, two-sided Wilcoxon signed-rank test, p = 0.393
- Figure S2c: Angular velocity CCW block: - 0.261 rad./sec., median vs. Angular velocity CW block: + 0.303 rad./sec., median, two-sided Wilcoxon signed-rank test, p < 0.001
- Figure S2e: reorientation bias ITI CCW block: - 0.011 ± 0.010 rad., mean ± s.e.m., Reorientation bias ITI CW block: - 0.016 ± 0.008 rad. two-sided Wilcoxon signed-rank test, p = 0.633
- Figure S2f: reorientation bias short ITIs: + 0.07 ± 0.029 rad., mean ± s.e.m. vs. reorientation bias shuffle short ITIs: + 0.0007 ± 0.028 rad., mean ± s.e.m.. reorientation bias long ITIs: + 0.0002 ± 0.053 rad., mean ± s.e.m., reorientation bias shuffle long ITIs: - 0.003 ± 0.004 rad., mean ± s.e.m.. reorientation bias short ITIs vs. reorientation bias shuffle short ITIs, two-sided Wilcoxon signed-rank test, p<0.05, reorientation bias long ITIs vs. reorientation bias shuffle long ITIs, two-sided Wilcoxon signed-rank test, p = 0.98
- Figure S3b: reorientation magnitude ctrl: + 1.363 ± 0.115 rad., mean ± s.e.m., reorientation magnitude ctrl shuffle: + 0.715 ± 0.044 rad., mean ± s.e.m., reorientation magnitude ablation: + 1.345 ± 0.129 rad., mean ± s.e.m., reorientation magnitude ablation: + 0.844 ± 0.122 rad., mean ± s.e.m.. ctrl vs. ctrl shuffle, two-sided Wilcoxon signed-rank test, p < 0.001, ablation vs. ablation shuffle, two-sided Wilcoxon signed-rank test, p < 0.01
- Figure S3c: swim rate control group: + 0.844 ± 0.027 swim/sec.; swim rate ablation group: + 0.822 ± 0.027 swim/sec., mean ± s.e.m.; two-sided T-test, p = 0.562
- Figure S3d: inter-swim interval control group: + 0.898 ± 0.021 sec.; inter-swim interval ablation group: + 0.940 ± 0.021 sec., mean ± s.e.m.; two-sided T-test, p = 0.164
- Figure S3e: distance traveled during swim control group: + 1.972 ± 0.047 mm.; distance traveled during swim ablation group: + 2.001 ± 0.049 swim/sec., mean ± s.e.m.; two-sided T-test, p = 0.668
- Figure S3f: swim duration control group: + 0.264 ± 0.007 sec.; swim duration ablation group: + 0.264 ± 0.007 swim/sec., mean ± s.e.m.; two-sided T-test, p = 0.932
- Figure S5d: percentage ON: + 34.5 %, median, OFF fraction: + 16.9 %, median. ON fraction vs. OFF fraction, two-sided Wilcoxon signed-rank test, p < 0.05
- Figure S7b: MI r-dHb: 0.26, median, MI iIPN: 1.08, median; r-dHb vs. iIPN, two-sided Mann-Whitney U test, p < 0.001
- Figure S7c: Bonferroni corrected two-sided Mann-Whitney U test, p < 0.05
- Figure S9d: neurons count dPPR^ipsi^: 24, median, neurons count dPPR^contra^: 29, median
- Figure S11a: ctrl: two-sided Wilcoxon signed-rank test, p = 0.570. treatment: two-sided Wilcoxon signed-rank test, p = 0.296
- Figure S11c: ctrl vs. treatment: two-sided Mann-Whitney U test, p < 0.299
- Figure S11e: ctrl: two-sided Wilcoxon signed-rank test, p = 0.687. treatment: two-sided Wilcoxon signed-rank test, p = 0.9
- Figure S11f: ctrl: two-sided Wilcoxon signed-rank test, p = 0.312. treatment: two-sided Wilcoxon signed-rank test, p = 0.688
- Figure S11h: ctrl: two-sided Wilcoxon signed-rank test, p = 0.305. treatment: two-sided Wilcoxon signed-rank test, p < 0.05

## Data and code availability

All the data described in the paper, the analysis notebooks and the figure generation notebooks will be made available upon publication.

## Supplementary figures

Eleven supplementary supplementary figures follow.

**Figure S1.**
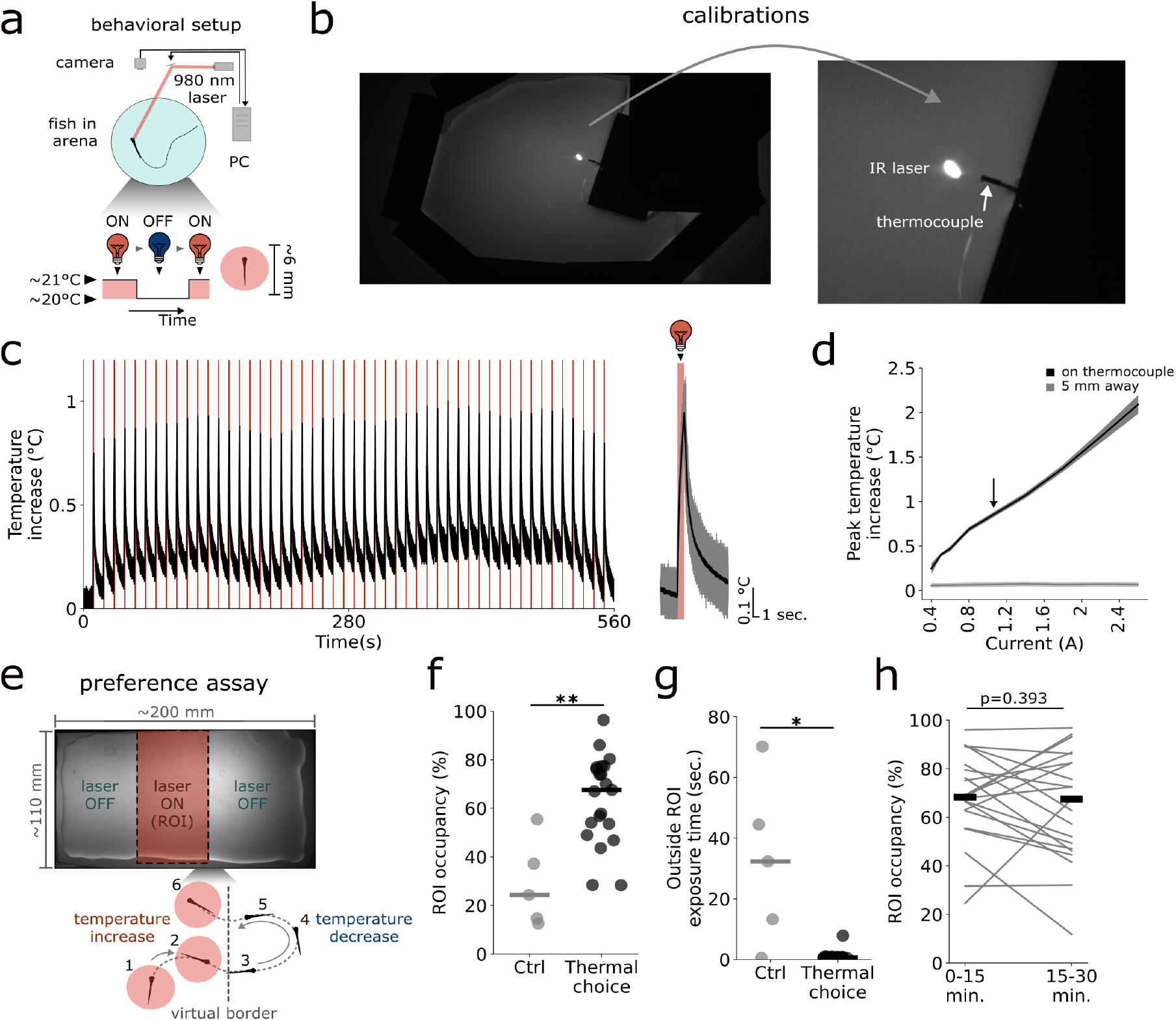
Cold avoidance with a laser-based setup. **(a)** I: Behavioral setup used for freely swimming experiments. **(b)** Image of the IR laser and the thermocouple during calibrations. **(c)** Left: example temperature recording during a calibration session. IR laser was directed to the thermocouple for 1 second, with 10 seconds interval. Right: stimulation triggered average. In black, average temperature increases and in gray single calibration trials. In red, laser ON period. **(d)** Calibration curve (mean ± s.e.m.). Laser power was selected based on these results (black arrow). Black: temperature increase recorded when the IR beam was directed on the sensor. Gray: temperature increase recorded when the IR beam was directed 5 millimeters away from the sensor. **(e)** Cold avoidance assay. Fish body temperature depends on its position in the arena. Within the ROI (a rectangular region at the center), the animal experiences a warmer temperature. **(f)** ROI occupancy in control (gray) and experimental condition (black). Each dot represent the performance of a single fish, the horizontal bar the median across animals. **(g)** Median time spent outside the ROI at each exit. Each dot represent the performance of a single fish, the horizontal bar the median across animals. **(h)** ROI occupancy during the first and second halves of the 30-minute experiment. Each line represents the performance of a single fish, the horizontal bar the median across animals.

**Figure S2.**
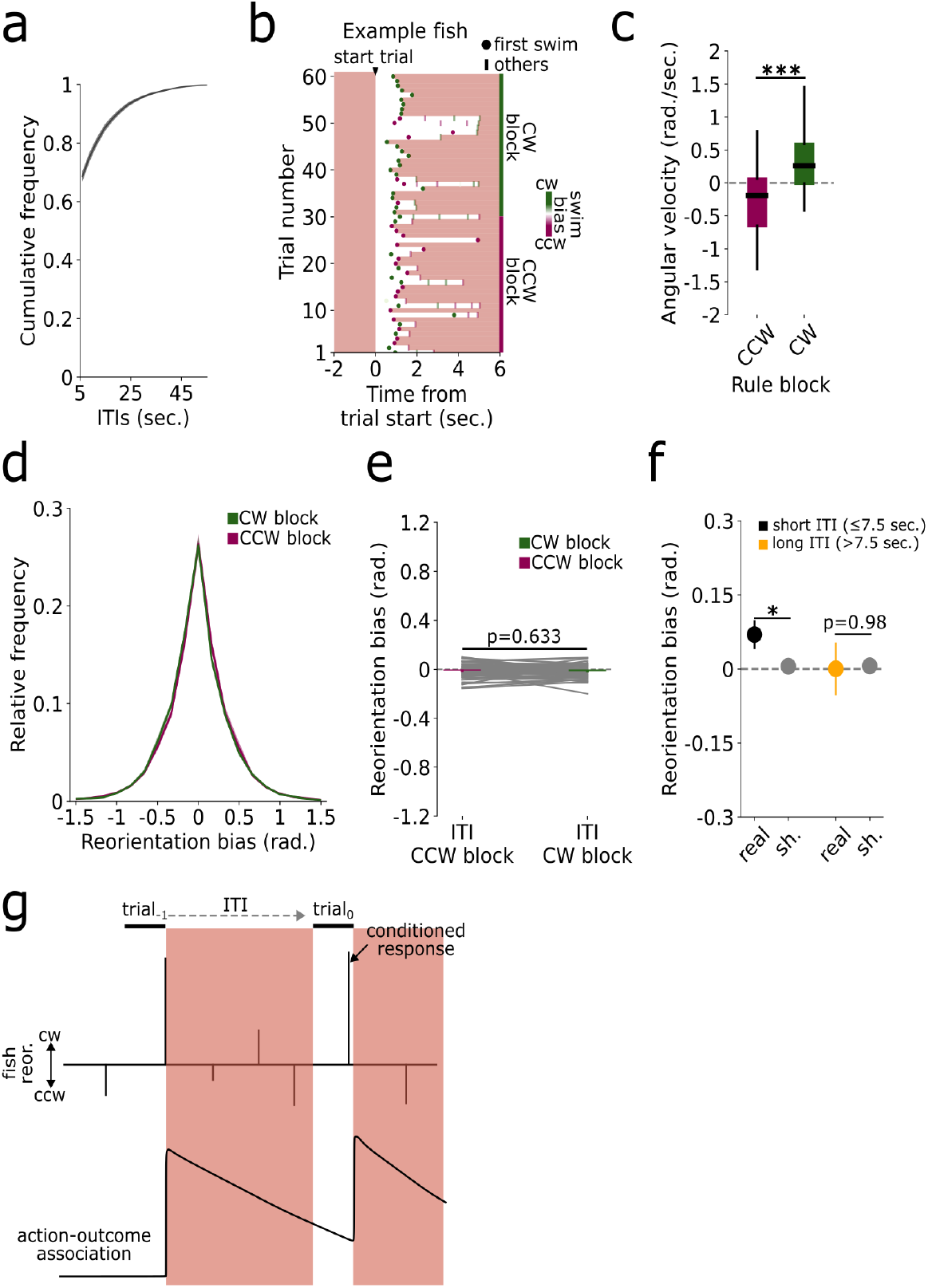
Operant thermoregulatory assay. **(a)** Cumulative distribution of ITIs during the operant assay (mean ± s.e.m.). **(b)** Example fish. Each dot represents the timing of the first swim, while bars represent subsequent swims. Color indicates turn direction (magenta: CCW, green: CW). **(c)** Average angular velocity from 0.5 to 0.9 seconds after temperature decrease (magenta: CCW block, green: CW block). **(d)** Relative frequency distribution of the reorientation bias (mean ± s.e.m.) during the ITIs of CCW (magenta) and CW (green) blocks. **(e)** Average reorientation bias (mean ± s.e.m.) during the ITIs of CCW (magenta) and CW (green) blocks. Each gray line represent a single fish. **(f)** Average reorientation bias (mean ± s.e.m.) evaluated after a short (black, ≤ 7.5 seconds) or a long (yellow, > 7.5 seconds) ITI (gray: ITI shuffles). **(g)** Schematic of the hypothesized process taking place during the operant assay. **(h)** Reaction time (mean ± s.e.m.). Each bar represents the time between temperature decrease and the first choice (black: 0 sec. delay, dark brown: 0.4 sec. delay, light brown: 0.8 sec. delay).

**Figure S3.**
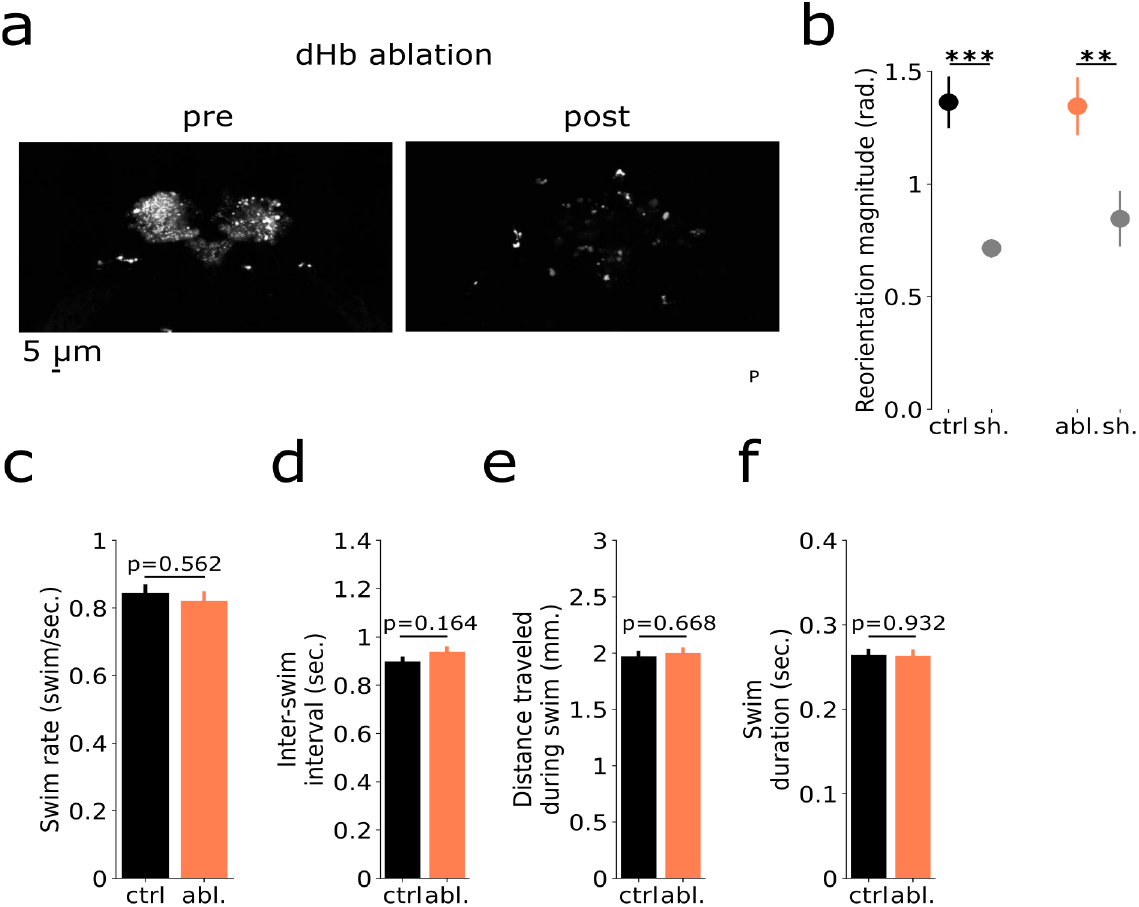
dHb chemogenetic ablation experiment. **(a)** Confocal image of a pre- (left) and post-dHb ablated fish (right). **(b)** Reorientation magnitude at temperature decrease stimulation for controls (black) and ablated group (pink) along with their shuffle controls. **(c)** Average swim rate (mean ± s.e.m.) for control (black) and ablation (pink) group. **(d)** Average inter-swim interval duration (mean ± s.e.m.) for control (black) and ablation (pink) group. **(e)** Average distance traveled during a swim (mean ± s.e.m.) for control (black) and ablation (pink) group. **(f)** Average swim duration (mean ± s.e.m.) for control (black) and ablation (pink) group.

**Figure S4.**
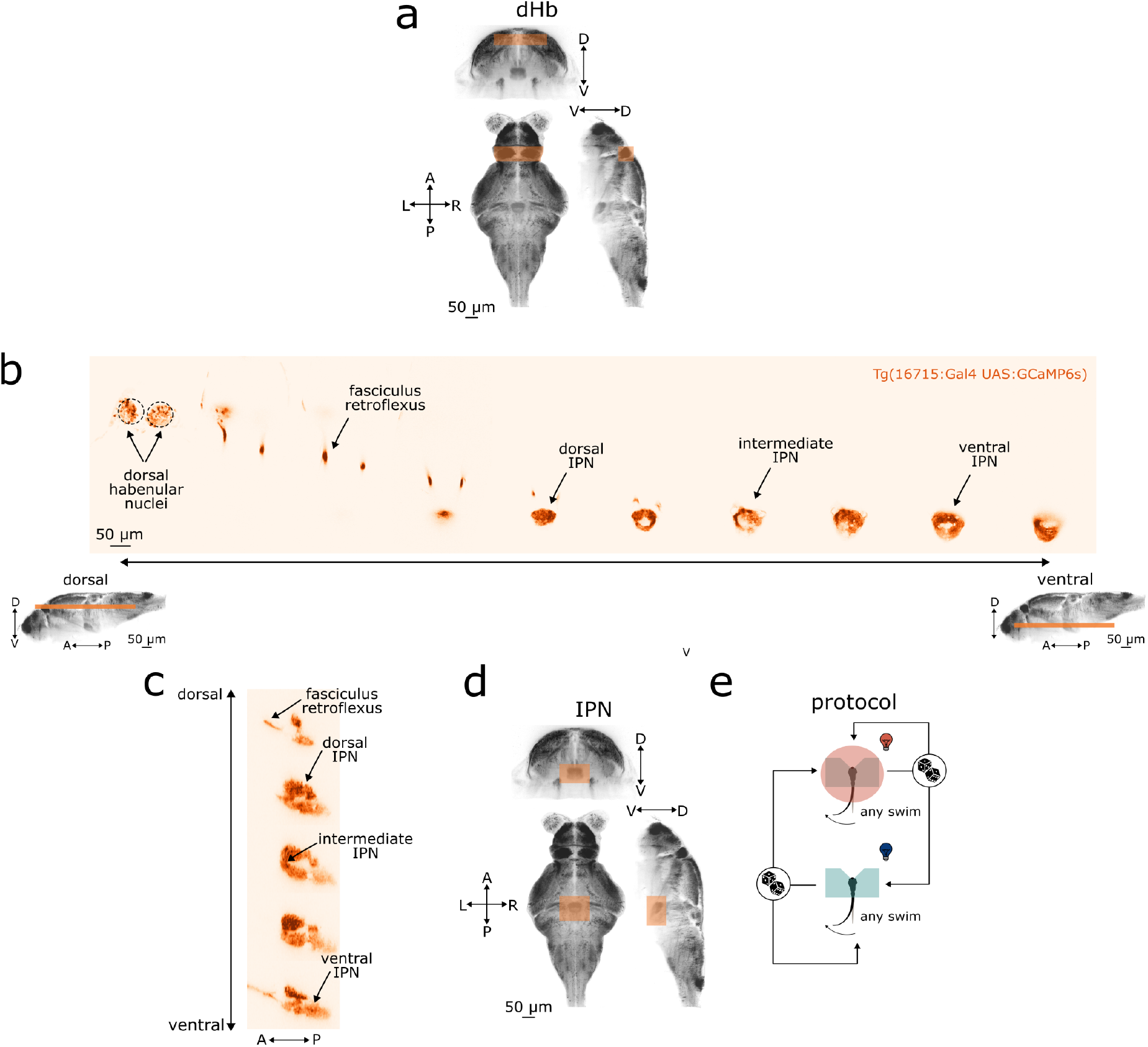
dHb-IPN pathway. **(a)** dHb anatomical position. Top: frontal projection. Left bottom: horizontal projection. Right bottom: sagittal projection. Reference space and anatomy from mapZebrain atlas. **(b)** 2-photon horizontal sections of dHb-to-IPN projections. **(c)** Sagittal sections of dHb axons in the IPN. **(d)** IPN anatomical position. Top: frontal projection. Left bottom: horizontal projection. Right bottom: sagittal projection. Reference space and anatomy from mapZebrain atlas. **(e)** Schematic of the head-fixed closed-loop protocol.

**Figure S5.**
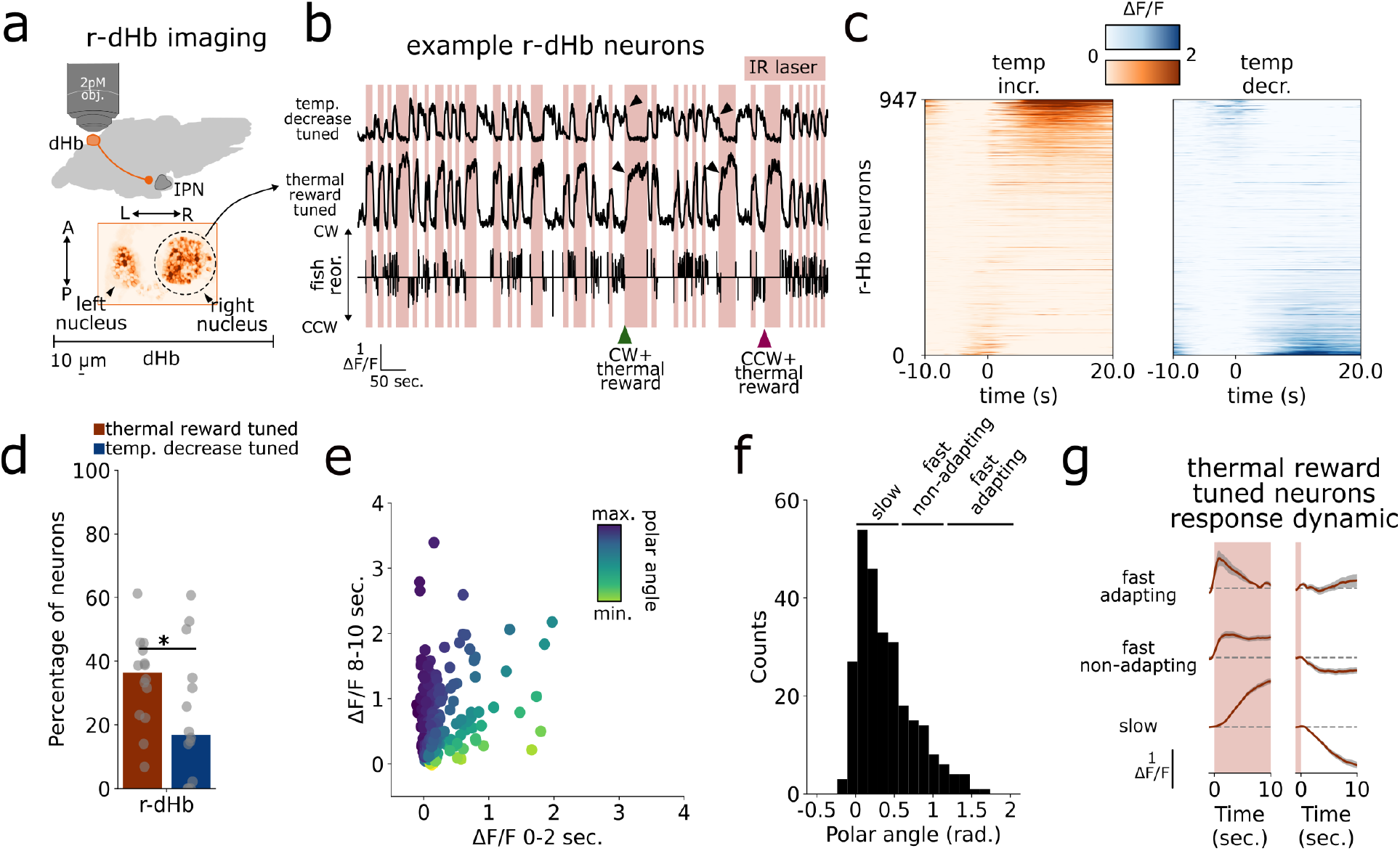
r-dHb calcium imaging experiment. **(a)** I: Top: Schematic of r-dHb 2-photon calcium imaging experiments. Bottom: Sum projection from an example recording. **(b)** Example of temperature decrease (top) and thermal reward (bottom) tuned neurons. The black trace below represents the animal’s behavior, with upward and downward spikes indicating CW and CCW reorientations, respectively. Green and magenta arrowheads point to CW+thermal reward and CCW+thermal reward events, respectively. **(c)** Triggered average at thermal reward and temperature decrease. Each row represents the activity of a single neuron, with correspondence between the left and the right plot. Traces have been sorted by peak activity at thermal reward. **(d)** Percentage of tuned neurons in the r-dHb. Each dot represents the percentage for a single fish, with bars indicating the median. **(e)** Polar angle computation. Each dot is a neuron, color-coded by the polar angle. **(f)** Polar angle distribution of r-dHb neurons. **(g)** ON neuron response dynamics (mean ± s.e.m.). Each line shows the average activity for response types (rows) across thermal reward/temperature decrease events (columns).

**Figure S6.**
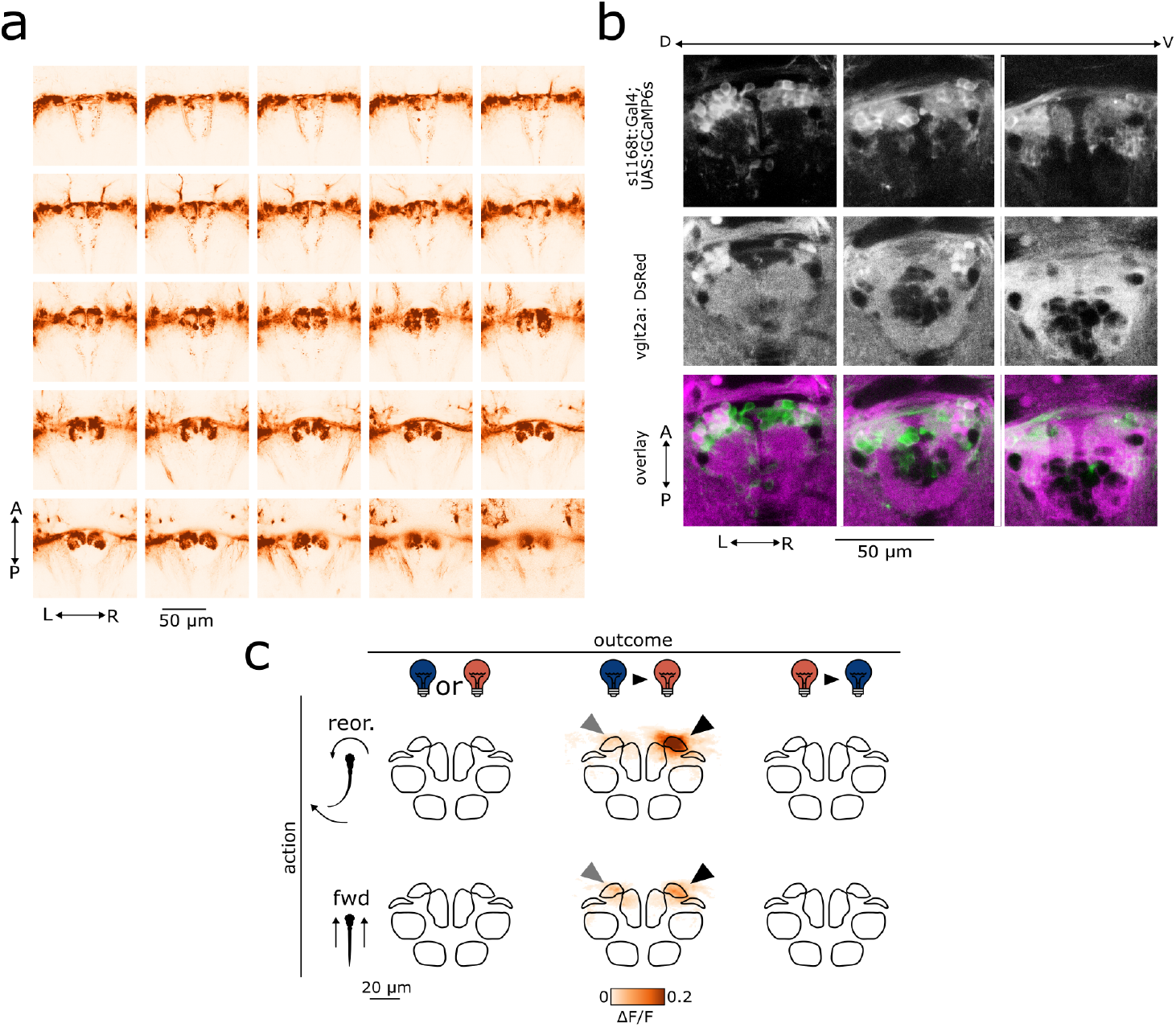
iIPN calcium imaging experiment. **(a)** 2-photon horizontal sections of the transgenic line labeling iIPN neurons. Dorsoventral planes are shown from top left to bottom right. **(b)** 2-photon horizontal sections where in red (appearing magenta in the image) are labeled excitatory processes and in green the imaged iIPN neurons with the corresponding glomeruli.**(c)** Average ΔF/F projection across action-outcome combinations. Rows represent actions, and columns represent outcomes. CCW and CW choices have been pooled together by flipping the stack along the midline. In this way, responses for the preferred direction can be seen on the right side, while responses to the opposite turn are on the left.

**Figure S7.**
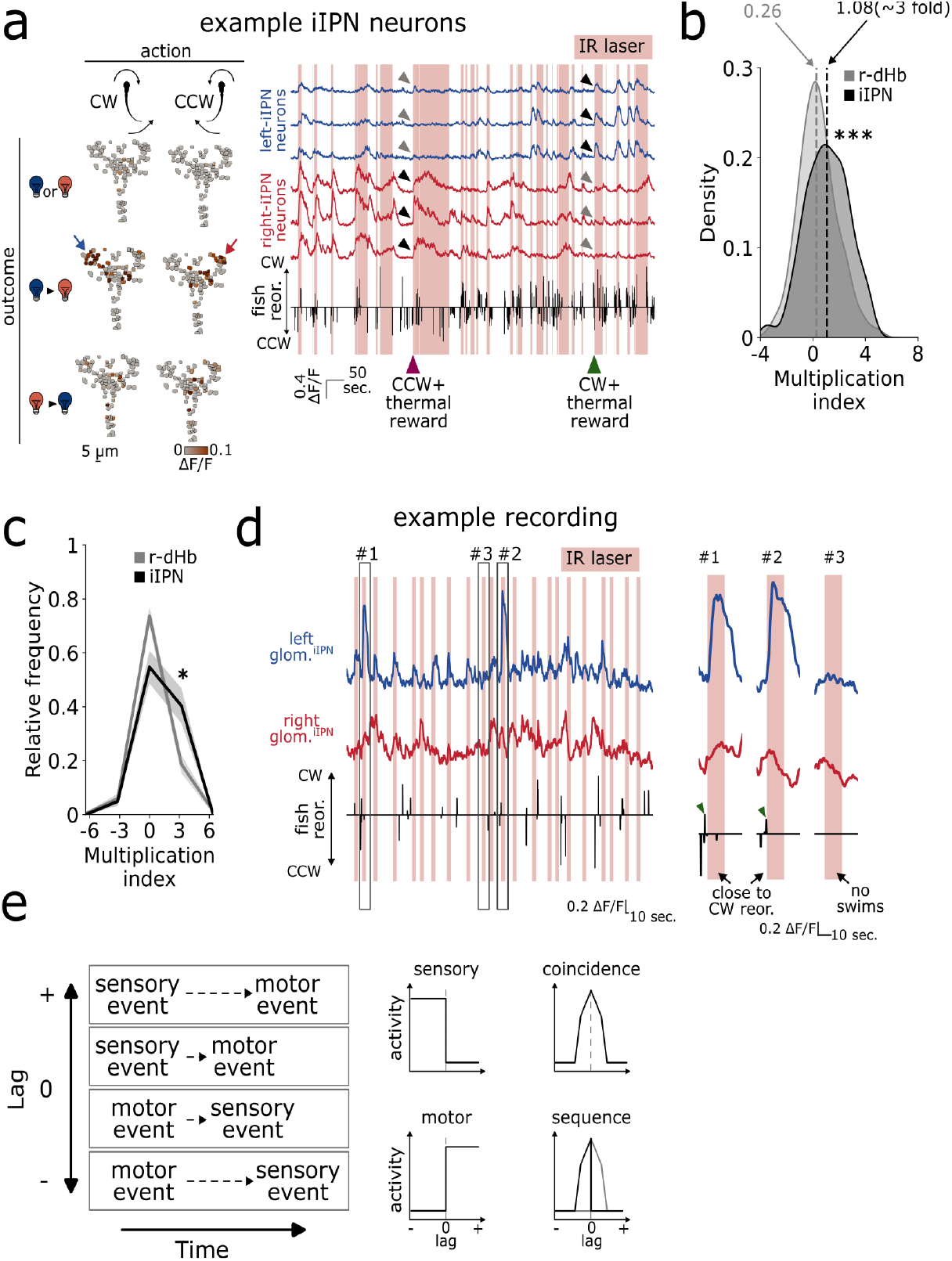
Somatic imaging of iIPN neurons and comparison with r-dHb coding. **(a)** Left: Average ΔF/F of an iIPN recording for CW and CCW turns (columns) and outcomes (rows). Right: Six example traces from the left (blue) and right (red) hemisphere. The black trace below represents the animal’s behavior, with upward and downward spikes indicating CW and CCW reorientations, respectively. Green and magenta arrowheads point to CW+thermal reward and CCW+thermal reward events, respectively. **(b)** Multiplication index density for r-dHb(gray) and iIPN(black) neurons. **(c)** Average relative distribution (mean ± s.e.m.) of the multiplication index for r-dHb (gray) and iIPN (black). **(d)** Left: Example recording from the left (top) and right (bottom) glomerulus during open-loop stimulation. Right: Magnified view of the events indicated by gray boxes in II (#1, #2, #3). **(e)** Left: Sensorimotor lag concept. Right: Possible outcomes for the sensorimotor lag analysis.

**Figure S8.**
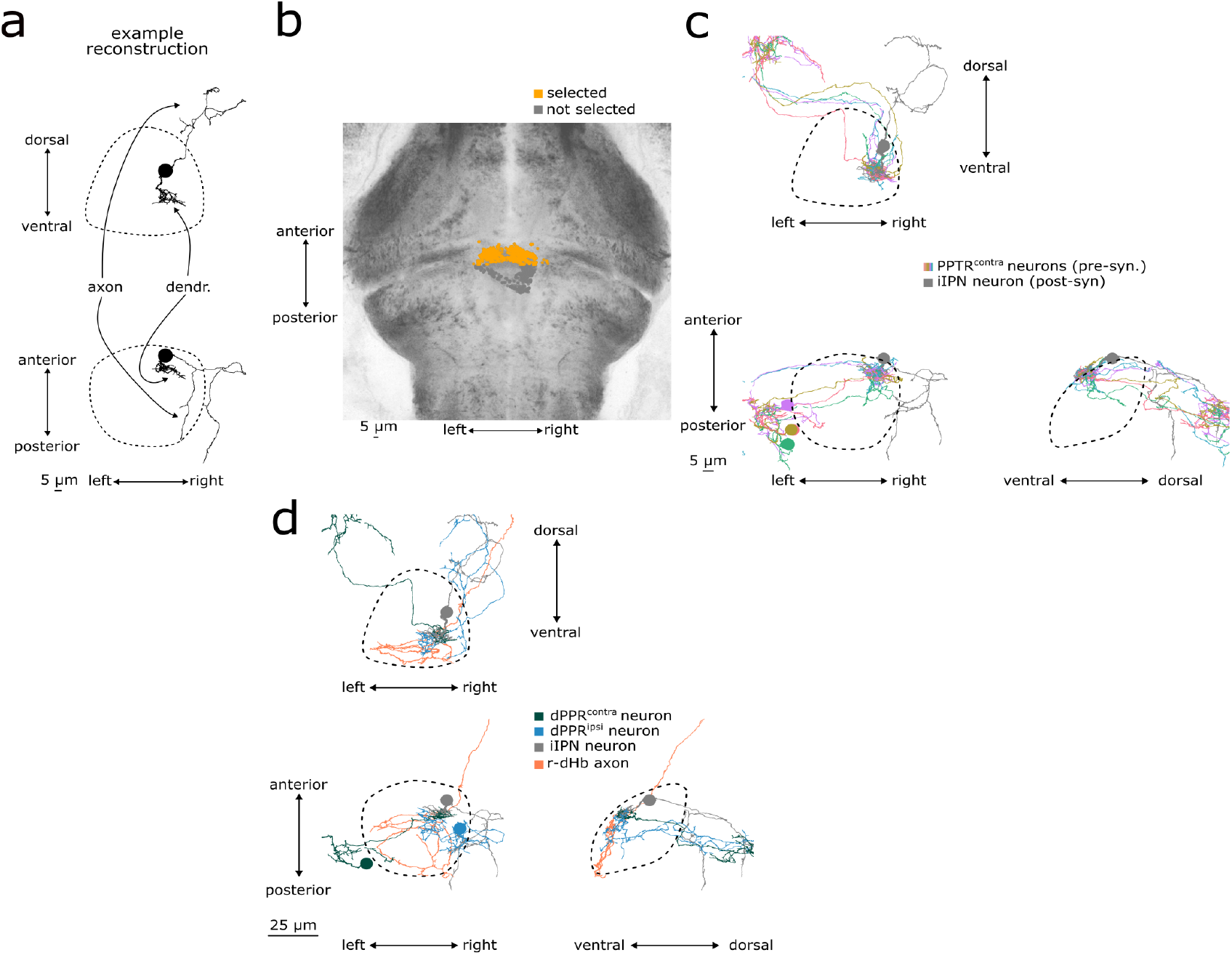
EM reconstructions. **(a)** Example iIPN neuron. Frontal projection on the top row and horizontal projection on the bottom row. Dashed regions enclose the IPN neuropil. **(b)** Detected contacts from dHb axons selected for the mediolateral distribution analysis (orange). Non-selected contacts are shown in gray. Horizontal section. Reference space and anatomy from mapZebrain atlas. **(c)** All reconstructed dPPR^contra^ neurons. An iIPN neuron is shown in gray. Colors for dPPR^contra^ neurons were randomly assigned. Top: frontal projection. Bottom left: horizontal projection. Bottom right: sagittal projection. Reference space from mapZebrain atlas. **(d)** Exemplar reconstructions: dPPR^ipsi^ axon projecting in the glomerulus (cyan), dPPR^contra^ axon targeting the dendrite of the glomerular iIPN neuron (green), r-dHb axon targeting the dendrite of the glomerular iIPN neuron (pink) and the iIPN neuron (gray). Bottom left: horizontal projection. Bottom right: sagittal projection. Dashed regions enclose the IPN neuropil. Reference space from mapZebrain atlas.

**Figure S9.**
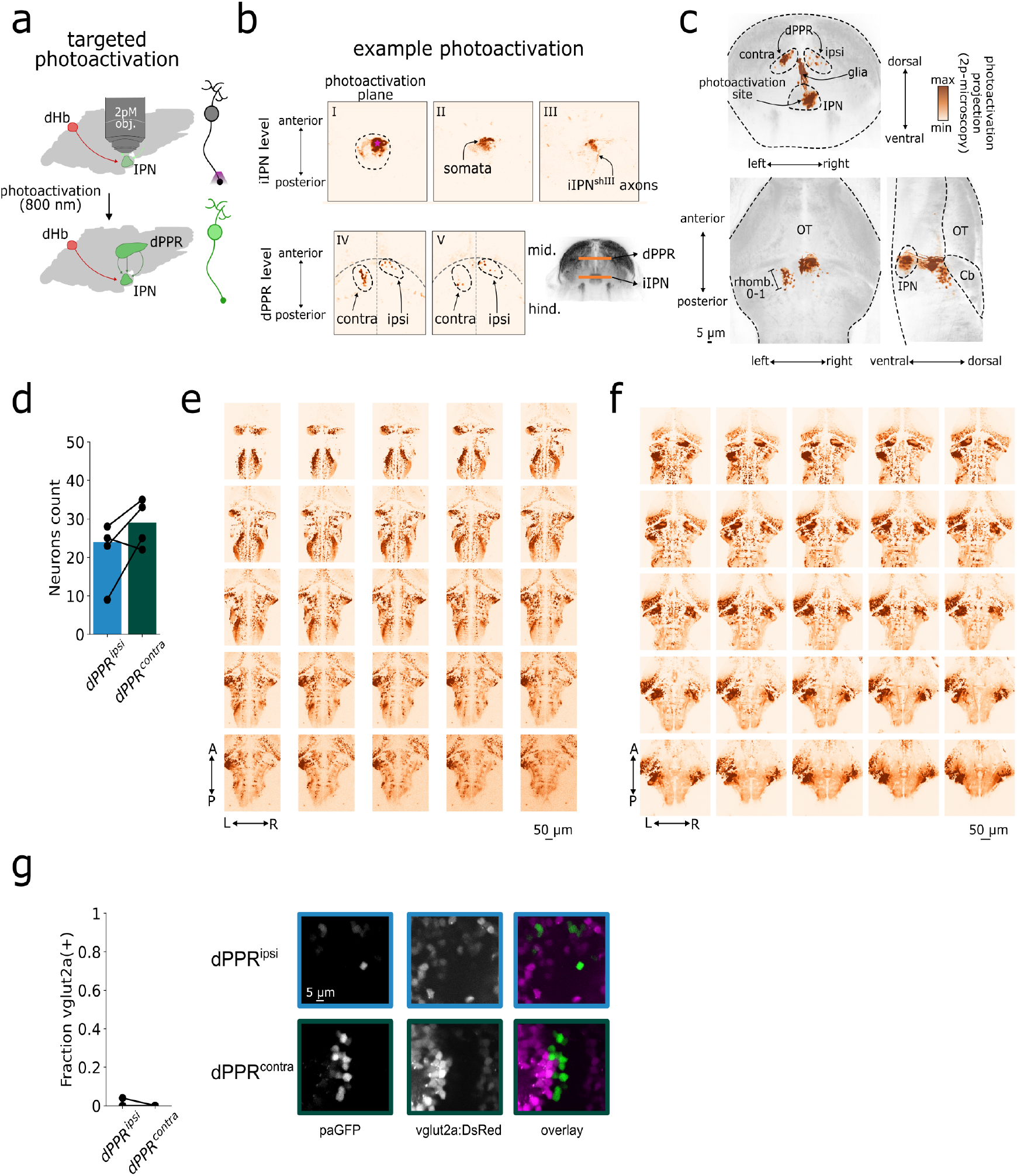
Targeted photoactivation experiment. **(a)** Schematic of targeted photoactivations. **(b)** Example photoactivation. Top row: photoactivation result in the iIPN. Magenta star indicates the scanned region. dHb axons, iIPN somata, and their axons are visible. Bottom row: photoactivation result in the dPPR. **(c)** Summary map of photoactivations. The map represents the average across fish, subtracted by control acquisitions where no photoactivation was performed. Top: frontal projection. Left bottom: horizontal projection. Right bottom: sagittal projection. **(d)** Cell count of labeled dPPR neurons in the ipsilateral (cyan bar) and contralateral (green bar) hemispheres, relative to the activated side. **(e)** 2-photon horizontal sections of the transgenic line labeling GABAergic neurons. Dorsoventral planes are shown from top left to bottom right. **(f)** 2-photon horizontal sections of the transgenic line labeling glutamatergic neurons. Dorsoventral planes are shown from top left to bottom right. **(f)** Left: Fraction of vglut2a positive dPPR^ipsi^ (cyan bar) and dPPR^contra^ (green bar) neurons. Right: example photoactivation experiment.

**Figure S10.**
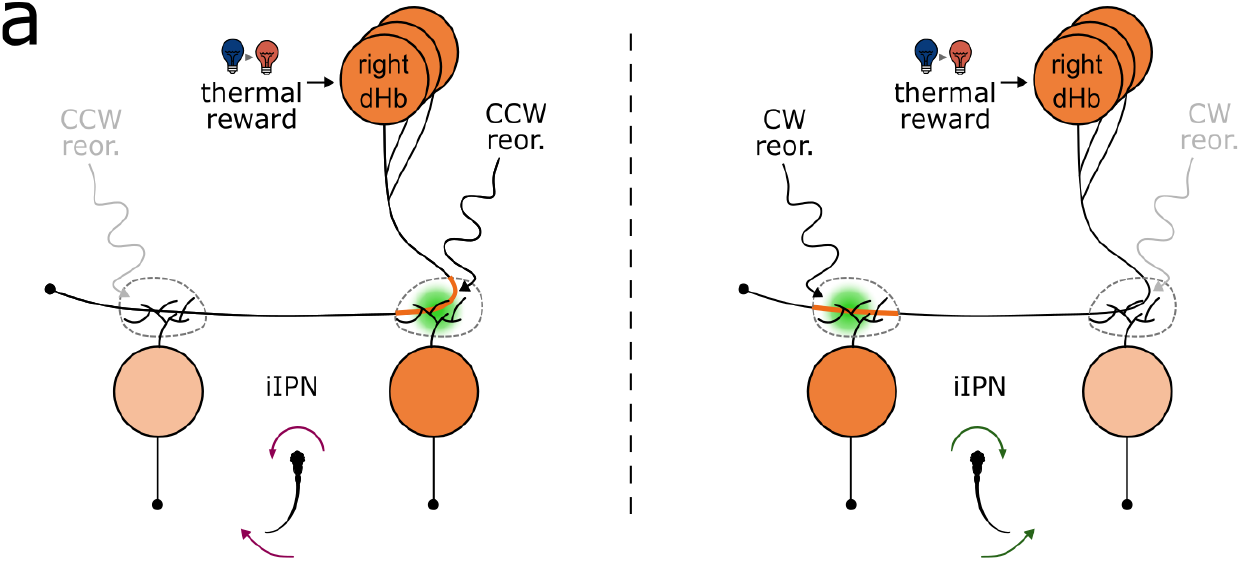
dHb axon modulation model. **(a)** Model schematic. dHb structure and function can be reconciled by modulation of axon terminals by a hypothetical neuronal population carrying reorientation information.

**Figure S11.**
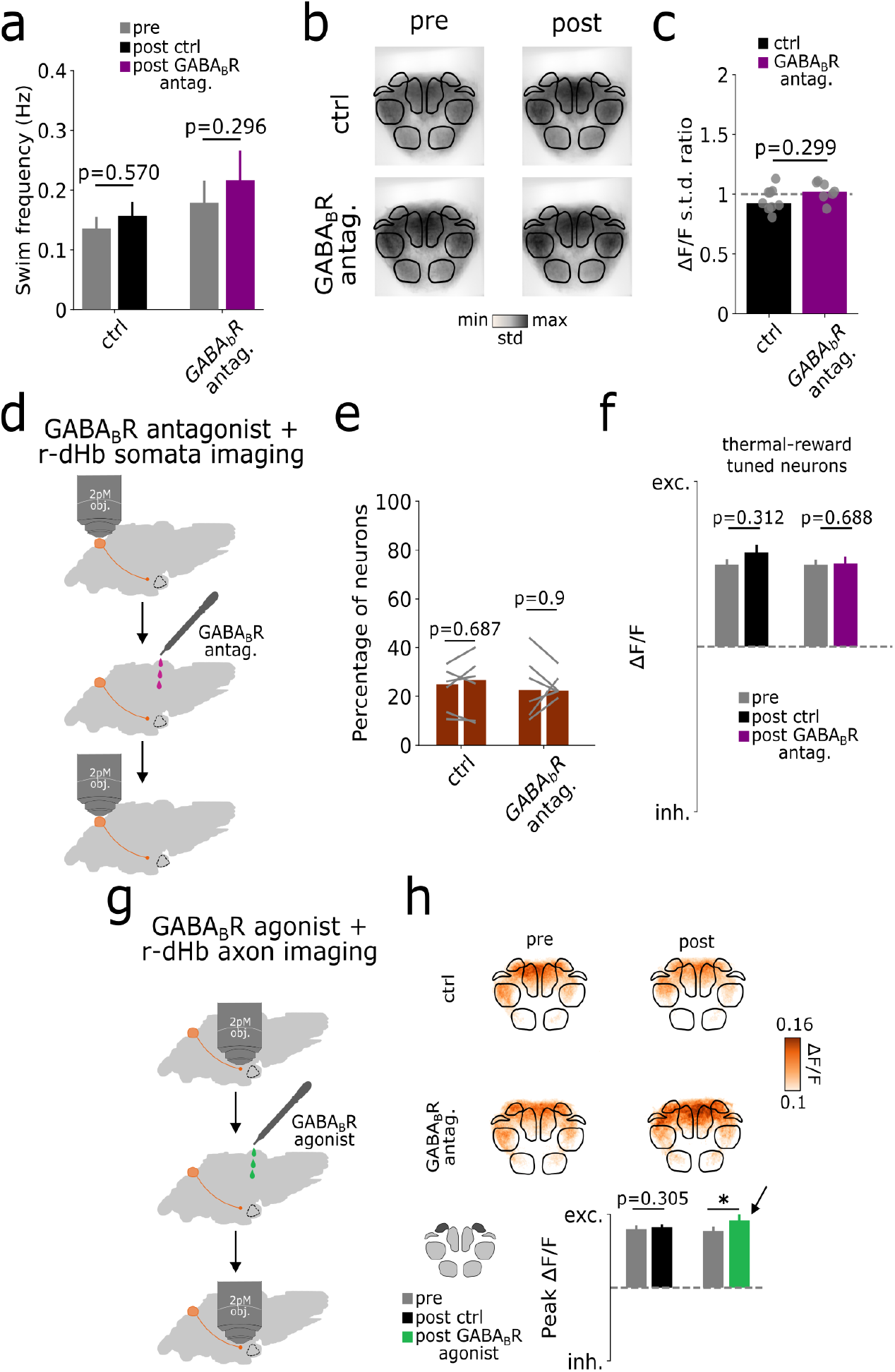
dHb axon pharmacology+imaging experiment. **(a)** Swim frequency (number of detected swims over the 30 minutes experiment) during the first imaging session (gray bars), during the second imaging session in the ctrl group (black bar) and during the second imaging session in the treatment group (purple bar). **(b)** Average pixel-wise ΔF/F standard deviation maps. Rows represent experimental groups (first row: ctrl group, second row: treatment group) and columns represent experimental timepoint (first column: first imaging session, second column: second imaging session). **(c)** Ratio between the average pixel-wise ΔF/F standard deviation map during the first and second imaging session. If close to 1, it implies that, on average, dHb axon are similarly active during the first and second session. **(d)** Schematic of GABA_B_R antagonist experiments in dHb somata. **(e)** Percentage of thermal reward tuned neurons in the r-dHb somata detected in the pharmacology experiments. **(f)** ΔF/F quantification (mean ± s.e.m.) for the thermal reward tuned neurons in the r-dHb somata (gray bars: first imaging session, black bar: second imaging session ctrl group, purples bars: second imaging session treatment group). **(g)** Schematicof GABA_B_R agonist experiments in dHb somata. **(h)** dHb axon activity upon GABA_B_R agonist treatment. Top: Peak ΔF/F projection during thermal reward. Rows represent experimental groups (first row: ctrl group, second row: treatment group) and columns represent experimental timepoint (first column: first imaging session, second column: second imaging session). Bottom: Peak ΔF/F quantification (mean ± s.e.m.) for different experimental groups. (gray bars: first imaging session, black bar: second imaging session ctrl group, purples bars: second imaging session treatment group).

